# Genome-scale CRISPR screening in a single mouse liver

**DOI:** 10.1101/2021.01.30.428976

**Authors:** Heather R. Keys, Kristin A. Knouse

## Abstract

A complete understanding of the genetic determinants underlying mammalian physiology and disease is limited by the availability of tools for high-throughput genetic dissection in the living organism. Genome-wide screening using CRISPR-Cas9 has emerged as a powerful method for uncovering the genetic regulation of cellular processes, but the need to stably deliver single guide RNAs to millions of cells has largely restricted its implementation to *ex vivo* systems. There thus remains a pressing need for accessible high-throughput functional genomics in a living organism. Here, we establish genome-wide enrichment and depletion screening in the liver of a single mouse and use this approach to uncover the complete regulation of cellular fitness in a living organism. We uncover regulation of cell fitness not previously identified in cell culture screens, underscoring the power of bringing high-throughput functional genomics into the organism. The approach we developed is accessible, robust, scalable, and adaptable to diverse phenotypes and CRISPR applications. We have hereby established a foundation for high-throughput functional genomics in a living mammal, enabling unprecedented insight into physiology and disease.

## INTRODUCTION

Our ability to understand and modulate mammalian physiology and disease requires the capacity to determine how all genes contribute to any given phenotype. In cell culture systems, genome-wide screening approaches have provided the power to identify all genes that positively or negatively regulate a cellular process in a single experiment. However, these *ex vivo* systems cannot reproduce all of the cellular processes that occur *in vivo* and, even when they can, they cannot recapitulate the entirety of extracellular factors that influence these phenomena *in vivo*. Although these limitations warrant studying cellular processes directly in the organism, probing complex phenotypes in the organism has historically required sacrificing experimental tractability. When it comes to inferring causal relationships, one is largely restricted to analyzing a single gene at a time using knockout mice. This discordance between experimental tractability and physiologic relevance has long limited our ability to understand mammalian physiology and disease. There is therefore a pressing need to bring high-throughput functional genomics into the organism.

Among the methods for high-throughput functional genomics in cell culture, genome-wide screening using CRISPR-Cas9 has emerged as a powerful approach^1, 2^. This entails delivering a single guide RNA (sgRNA) library alongside Cas9 to cells, an intervening period for protein depletion and phenotypic selection, and ultimately deep sequencing to detect changes in sgRNA abundance and thus hits in the screen. Critical to the success of the screen is delivering sgRNAs in a manner that is stable and has high coverage, with any given sgRNA being delivered to at least a few hundred cells, so that changes in sgRNA abundance can be evaluated at the end of screening with the power to identify significantly enriched and depleted sgRNAs. Lentivirus is the preferred method for delivering sgRNAs into cells as it allows for stable, single-copy integration of an sgRNA within a cell. While lentivirus is effective at delivering genome-wide sgRNA libraries to tens of millions of cells in culture, it is less efficient at delivering sgRNAs to cells *in vivo*. Although groups have successfully used lentivirus to transduce keratinocytes and neurons *in vivo*, the number of cells transduced per mouse is limited and previous approaches required over 50 mice to be pooled for a single genome-scale screen^3, 4^. As such, genome-wide CRISPR screening has largely been limited to cell culture systems or cellular transplantation models^5, 6^.

Bringing genome-scale screening into the organism requires overcoming the barriers to stable, high-coverage sgRNA delivery into tissues. In this regard, the mouse liver is an appealing target organ. Comprised of tens of millions of hepatocytes, a single mouse liver offers cell numbers compatible with genome-scale screening. Moreover, given the liver’s diverse metabolic functions and impressive regenerative capacity, hepatocytes exhibit a broad range of phenotypes that are ripe for genetic dissection. Unfortunately, efforts to deliver lentivirus to the liver have suffered from poor transduction efficiency and immune-mediated clearance of transduced hepatocytes^7, 8^. Although groups have leveraged other delivery methods to perform genetic screens in the liver, these methods all have limitations that prevent genome-scale, enrichment and depletion screening. For example, adeno-associated virus (AAV) can be used to deliver sgRNAs to the majority of hepatocytes, but because AAV vectors typically remain episomal the sgRNAs will not persist in proliferating cells. Analysis of AAV-based screens therefore requires individually amplifying and sequencing the genes targeted by the sgRNAs, dramatically limiting the number of genes that can be screened in any given experiment^9^. Hydrodynamic tail vein injection of plasmids encoding transposons can also introduce sgRNAs into hepatocytes and offers the benefit of integration into the host genome. However, hydrodynamic tail vein injection only transfects 10-40% of hepatocytes in a non-uniform manner, again limiting the scale of the screen^10–13^. As such, genetic screens in the liver have largely been restricted to small-scale screens on the order of tens to hundreds of genes.

Here, we have established accessible, genome-wide, enrichment and depletion screening in the mouse liver. We developed an approach for stable, high-coverage delivery of a genome-scale sgRNA library into the liver of inducible Cas9 mice, allowing for Cas9 induction and phenotypic selection at any point in the animal’s lifetime. To validate this approach, we performed a screen for hepatocyte fitness in the neonatal liver. Our screen had the ability to uncover positive and negative regulators of hepatocyte fitness in individual mice with high reproducibility across mice. We discovered genes with sex-specific effects on hepatocyte fitness as well as genes that are uniquely required for hepatocyte fitness in a living organism. This approach is accessible and adaptable to diverse phenotypes and CRISPR methods and thereby provides a foundation for high-throughput genetic dissection in a living organism.

## RESULTS

### Genome-scale sgRNA delivery in a single mouse liver

Given the advantages of lentiviral-mediated sgRNA delivery over other sgRNA delivery methods, we sought to establish efficient lentiviral sgRNA delivery to the liver. We hypothesized that intravenously injecting highly concentrated lentivirus into neonatal mice might avoid the poor transduction efficiency and immune clearance observed by others^14^. To test this, we generated lentiviruses encoding a non-targeting sgRNA (sgAAVS1) alongside mCherry or mTurq2 reporters and injected varying doses of an equal mixture of these two lentiviruses into postnatal day (PD) one mice (Figure 1A). We observed a dose-dependent increase in the percentage of transduced hepatocytes, with a dose of 5 x 10^7^ transduction units (TU) transducing over 75% of hepatocytes (Figures 1B and 1C). Importantly, these transduced hepatocytes were distributed uniformly throughout the liver lobule and persisted into adulthood (Figures 1B, S1A, and S1B). Based on measurements of hepatocyte and liver volume and the transduction frequency of two fluorescent reporters, we estimated that a dose of 5 x 10^7^ TU transduces approximately 10 million hepatocytes per PD1 liver with an average of just two integration events per cell (Figures 1C and S1C). This transduction efficiency would afford >200-fold coverage of a 100,000 feature sgRNA library in a single mouse. Our lentiviral approach therefore establishes an sgRNA delivery method that is wholly compatible with genome-scale screening in the mouse liver.

**Figure 1.**
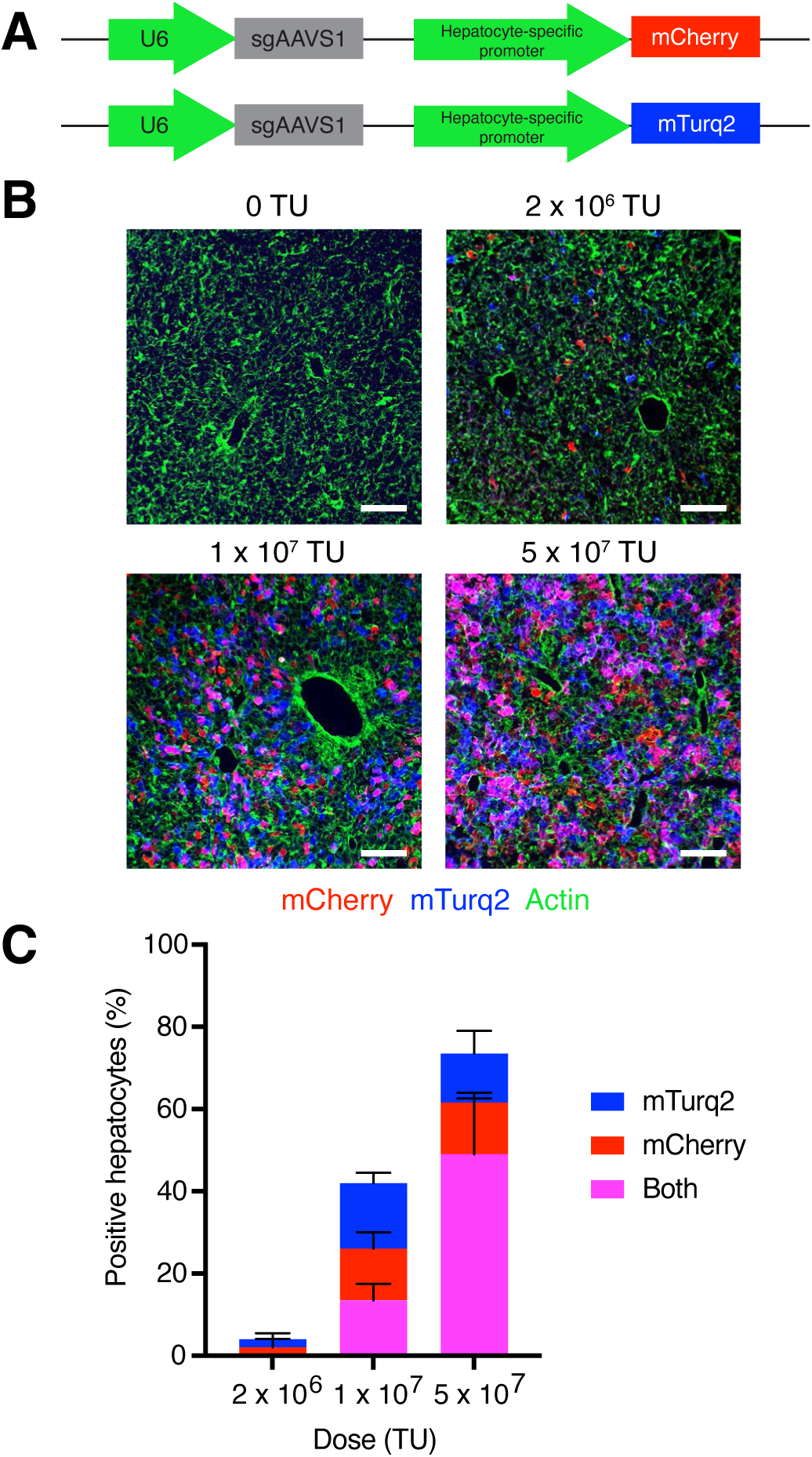
Genome-scale sgRNA delivery in a single mouse liver. (A) Lentiviral vectors for U6-driven expression of an sgRNA and hepatocyte specific-expression of a fluorescent reporter (mCherry or mTurq2). (B) Images of endogenous mCherry and mTurq2 fluorescence in livers from mice four days after injection with an equal mixture of sgAAVS1-mCherry and sgAAVS1-mTurq2 lentiviruses. Livers were counterstained with phalloidin (green) to label actin. Scale bars, 100 µm. (C) Percent mCherry-, mTurq2-, and double-positive hepatocytes in livers from mice four days after injection with an equal mixture of sgAAVS1-mCherry and sgAAVS1-mTurq2 lentiviruses. Error bars indicate standard deviation. n = 3 mice per dose and 200 hepatocytes per mouse. See also Figure S1.

We next asked whether we could use this sgRNA delivery approach as the basis for temporally-controlled protein depletion in hepatocytes. We used commercially available loxP-stop-loxP-Cas9 (LSL-Cas9) mice in which we could induce Cas9 in nearly all hepatocytes by injecting an adeno-associated virus expressing Cre recombinase from the hepatocyte-specific *Tbg* promoter^15^ (AAV-Cre, Figures S2A and S2B). To evaluate the kinetics and efficiency of protein depletion, we selected two long-lived, non-essential proteins: the mitochondrial enzyme MAO-B (encoded by *Maob*) and the nuclear lamin Lamin B2 (encoded by *Lmnb2*)^16, 17^. After delivering sgMaob-mCherry or sgLmnb2-mCherry lentivirus to PD1 mice, we injected PBS or AAV-Cre at PD5 and harvested livers at various time points to evaluate protein levels in individual hepatocytes (Figure 2A). By two weeks after Cas9 induction, MAO-B and Lamin B2 were depleted exclusively in mCherry-positive hepatocytes in mice injected with AAV-Cre (Figures 2B, 2C, S2C, and S2D). Importantly, this combination of lentiviral-mediated sgRNA delivery, AAV-Cre-mediated induction of Cas9, and resulting gene targeting did not induce detectable hepatocyte damage or liver inflammation (Figures S2E-G). Hepatocyte turnover was also unaffected (Figures S2H and S2I). This approach therefore offers an effective platform for hepatocyte-specific protein depletion and genetic screening at any point in the animal’s lifetime without detectable confounding perturbation of the cells or tissue.

**Figure 2.**
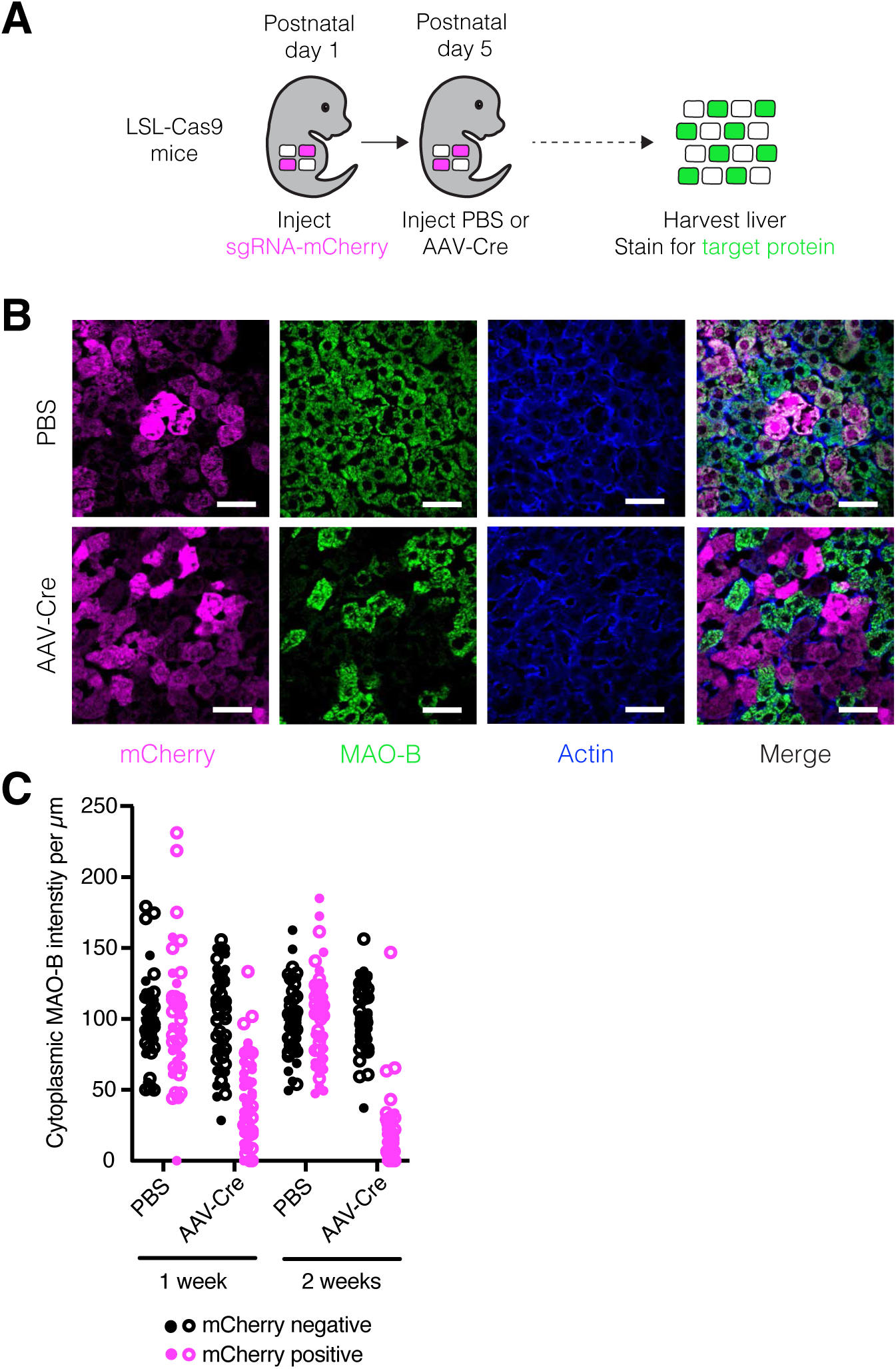
Temporally-controlled protein depletion in the mouse liver. (A) Scheme for inducing protein depletion in LSL-Cas9 mice. (B) Images of livers from LSL-Cas9 mice injected with sgMaob-mCherry followed by PBS or AAV-Cre immunostained for mCherry (magenta), MAO-B (green), and actin (blue). Scale bars, 45 µm. (C) Cytoplasmic MAO-B intensity per µm in mCherry-positive and mCherry-negative hepatocytes from LSL-Cas9 mice injected with sgMaob-mCherry followed by PBS or AAV-Cre. Closed and open circles represent values from male and female mouse, respectively. n = 1 male and 1 female mouse per condition and 25 cells per mouse. See also Figure S2.

### A genome-scale screen in the liver

To enable genome-scale screening for diverse hepatocyte phenotypes, we sought to prepare an sgRNA library targeting all genes expressed in the developing, quiescent, and regenerating mouse liver. We performed RNA sequencing on livers at various time points during mouse development and after liver injury and determined that 13,266 protein-coding genes were expressed (FPKM > 0.3) at one or more time points (Figures 3A, S3A, and S3B; Table S1). We generated an sgRNA library targeting 13,189 of these genes (average of 5 sgRNAs per gene) alongside a previously published set of 6,500 control sgRNAs (∼2,000 non-targeting sgRNAs and ∼4,500 sgRNAs targeting exonic and intronic regions of control genes)^18^ for a total of 71,878 unique sgRNAs (Figure S3C; Table S2).

**Figure 3.**
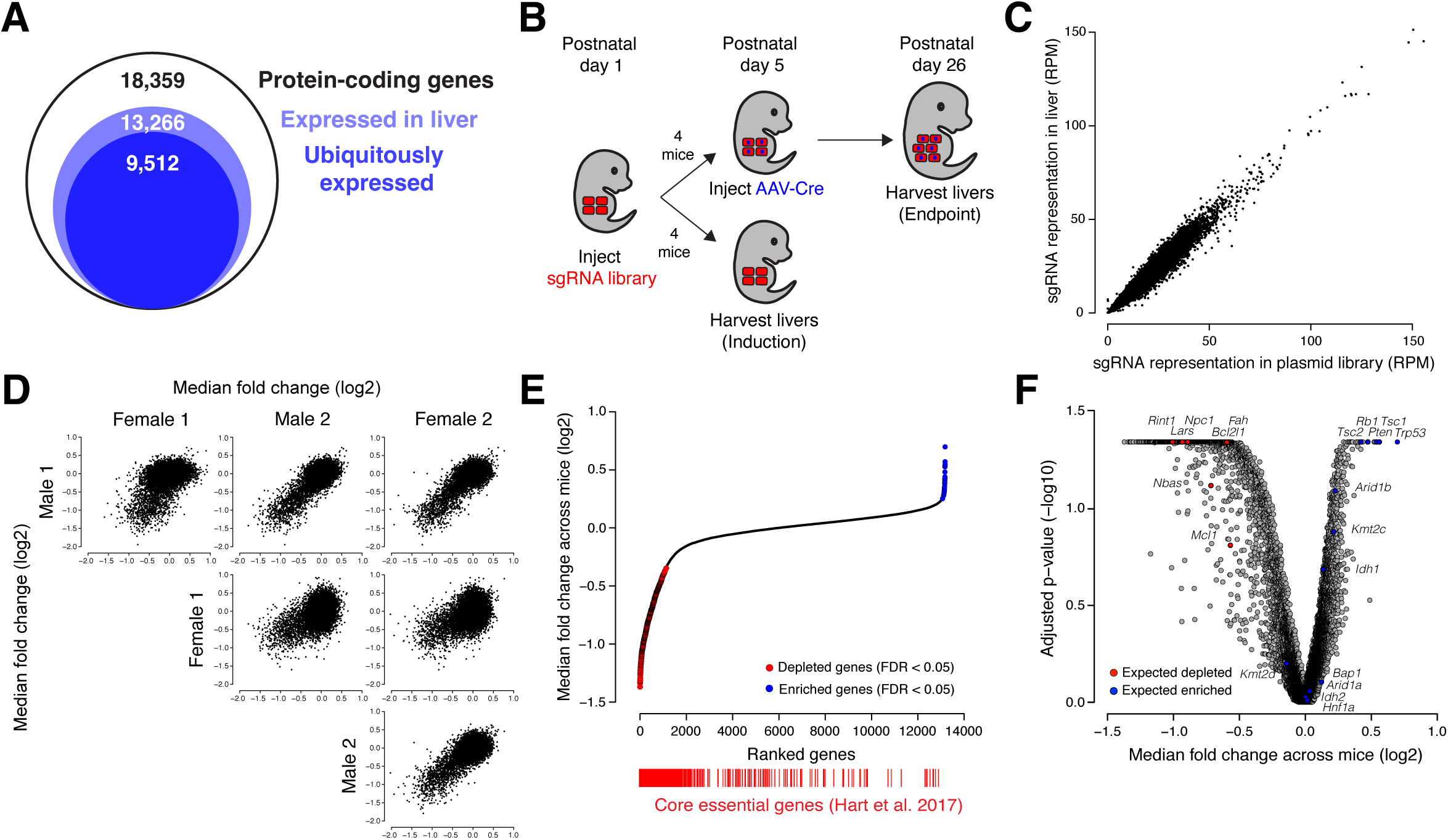
A genome-scale screen for hepatocyte fitness in the neonatal mouse liver. (A) Number of protein-coding genes expressed in the liver as determined by RNA sequencing of livers at various time points. (B) Scheme for performing a genome-scale screen for hepatocyte fitness in neonatal mice. (C) Representation of sgRNAs in livers four days after injection with lentiviral library relative to the sgRNA representation in the plasmid library expressed as reads per million (RPM). n = 2 male and 2 female mice pooled into a single sequencing library. Pearson correlation r = 0.97. (D) Pairwise comparisons of median fold change (log2) for each gene for each mouse at the endpoint of the screen. (E) Genes ranked by median fold change across mice (log2) with significantly depleted genes denoted by red points and significantly enriched genes denoted by blue points (FDR < 0.05 by two-tailed Wilcoxon test). Core essential genes (red bars) are positioned below based on gene rank to demonstrate their significant depletion across mice. p < 2.2 x 10^-16^ by one-sided Kolmogorov-Smirnov test. (F) Benjamini-Hochberg-adjusted Wilcoxon p-value (-log10) versus the median fold change across mice (log2) for each gene in the screen. Highlighted are control gene sets consisting of tumor suppressor genes in hepatocellular carcinoma (expected to enrich, blue) and genes required for hepatocyte viability (expected to deplete, red). Expected depleted p = 3.5 x 10^-6^ by one-sided Kolmogorov-Smirnov test, expected enriched p = 4.7 x 10^-4^ by one-sided Kolmogorov-Smirnov test. See also Figure S3.

With this method in hand, we undertook a genome-wide screen for hepatocyte fitness (Figure 3B). To screen for the ability of hepatocytes to both persist and proliferate, we elected to screen over a three-week period in neonatal development when hepatocytes undergo approximately three population doublings to increase liver mass. We injected 5 x 10^7^ TU of our lentiviral library into four female and four male LSL-Cas9 mice at PD1. At PD5, we harvested livers from two males and two females to evaluate the initial library representation. The sgRNA representation in these four livers correlated extremely well (Pearson r = 0.97) with the plasmid library and we detected all sgRNAs, indicating that we can effectively deliver and recover a genome-scale sgRNA library from the neonatal mouse liver (Figure 3C; Table S3). In the remaining mice, we induced Cas9 at PD5 and harvested their livers at PD26 to evaluate the final library representation.

We used the Model-based Analysis of Genome-wide CRISPR/Cas9 Knockout (MAGeCK) algorithm^19^ to identify enriched and depleted genes based on statistical differences in their change in sgRNA abundance at PD26 relative to PD5 (Table S3). Using a false discovery rate cutoff of 0.05, we identified 6, 0, 0, and 2 significantly enriched and 364, 40, 386, and 297 significantly depleted genes in male 1, female 1, male 2, and female 2, respectively, indicating that our method can detect enriched and depleted genes in a single mouse. We also generated gene-level scores by calculating the median log2 fold change in abundance of sgRNAs targeting a given gene. We observed a strong correlation in gene scores across the four mice (Pearson r = 0.46 to 0.75, Figures 3D and S3D; Table S3). Indeed, the reproducibility we observed across mice is similar to that observed in cell culture screens in cases where replicates are performed (Pearson r = 0.59 to 0.65)^20, 21^. We note that while Female 1 was less well-correlated with the rest of the mice, the correlation between Female 1 and other mice was still significant (p < 2.2 x 10^-16^) and we have no technical reason to invalidate Female 1’s data. We therefore considered all four mice as biological replicates without normalizing gene scores between mice for subsequent analyses.

To improve our power to identify significantly enriched and depleted genes, we combined the data from all four mice and calculated a unified gene score representing the median log2 fold change for each gene across mice (Table S3). Using a false discovery rate cutoff of 0.05, we identified 30 significantly enriched genes and 661 significantly depleted genes across all mice (Figure 3E, Table S3). Importantly, these gene scores were not positively correlated with gene expression level or protein half-life^16^, reaffirming that long-lived proteins were effectively depleted (Figures S3E and S3F). Collectively, these initial screen results establish the technical feasibility of genome-wide screening in the mouse liver. Importantly, while screening multiple mice in parallel increases the power to discover significant hits, a single mouse is sufficient to identify significantly enriched and depleted genes.

We next asked whether our screen reliably uncovered regulators of cell fitness. We first assessed whether genes established to be essential in cell culture^22^ were significantly enriched among the depleted genes. This set of core essential genes was indeed significantly depleted within each individual mouse and across all four mice (Figure 3E and S3G). To evaluate whether our screen could reliably reveal regulation specific to hepatocyte fitness in the liver, we assessed the gene scores for two sets of genes known to affect hepatocyte fitness in the organism: (1) a set of 13 genes established as tumor suppressors in hepatocellular carcinoma^23^ (expected to enrich) and (2) a set of seven genes required for hepatocyte viability (expected to deplete) (Table S3). Among the tumor suppressor genes, eight of the 13 genes were significantly enriched (FDR < 0.25, Figure 3F). Among the genes required for hepatocyte viability, all seven genes were significantly depleted (FDR < 0.25, Figure 3F). These results support our screen as a reliable platform for uncovering the genetic regulation of the phenotype in question.

### Genome-scale screening in the organism affords unique insights

Genome-scale screening in the organism, by virtue of preserving the native state and context of the cell and phenotype under investigation, enables several biological insights not possible in cell culture. One such advantage is the ability to screen wild-type cells that do not carry pre-existing mutations. Most cell lines naturally harbor or inevitably acquire mutations that improve their viability and proliferation in culture^24^, compromising the ability to query the function of these mutated genes in screens. As such, fitness screens in cell culture have an impaired ability to uncover tumor suppressor genes^25–28^. In contrast, our screen readily recovered tumor suppressor genes. Over half of the 25 most enriched genes in our screen are established to act as tumor suppressor genes in at least one context^29, 30^ (Table S3). Indeed, a set of the top 50 computationally predicted pan-cancer tumor suppressor genes^29^ was significantly enriched in our screen, but not in fitness screens in mouse embryonic stem cell lines^25, 26^ or human hepatocellular carcinoma cell lines^27, 28^ (Figures 4A and S4A; Table S3). Our screen’s unique ability to, within only a few population doublings, reliably recover tumor suppressors highlights the power of being able to screen unmutated, wild-type cells and thereby probe all genes that may influence a phenotype

**Figure 4.**
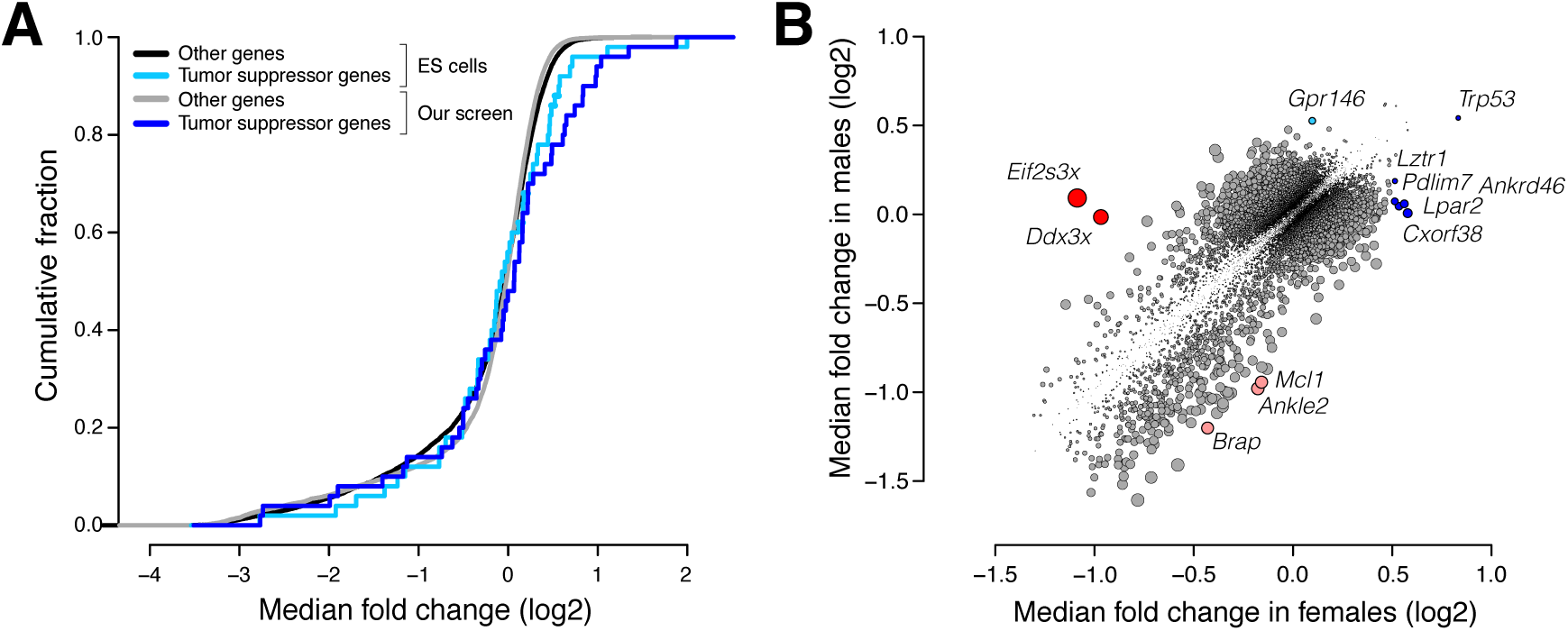
Genome-scale screening in the organism enhances discovery of tumor suppressor genes and uncovers genes with sex-specific effects. (A) Cumulative fraction of tumor suppressor genes (cyan, blue) and other genes (black, grey) based on quantile-normalized median fold change (log2) of their gene scores across screens in mouse embryonic stem (ES) cells and our screen. ES cells p > 0.05 by one-sided Kolmogorov-Smirnov test, our screen p = 6.5 x 10^-4^ by one-sided Kolmogorov-Smirnov test. (B) Median fold change (log2) across males versus median fold change (log2) across females for each gene. Highlighted are genes uniquely enriched in females (blue), genes uniquely enriched in males (cyan), genes uniquely depleted in females (red), and genes uniquely depleted in males (pink). Point size is proportional to the absolute difference in median log2 fold change between females and males. See also Figure S4.

A second advantage of screening directly in the organism is the opportunity to investigate how biological sex influences the relationship between genotype and phenotype. To determine whether biological sex affected hits in our screen, we compared the gene scores for all genes in males versus females. We identified three X-linked genes and nine autosomal genes with sex-specific effects on fitness (Figure 4B). The genes with the greatest difference between the sexes were the X-linked genes *Ddx3x* and *Eif2s3x* which were exclusively essential in females. Both *Ddx3x* and *Eif2s3x* facilitate protein synthesis. Both of these genes escape X inactivation and have paralogs on the Y chromosome, *Ddx3y* and *Eif2s3y*, with similar function^31, 32^. Thus, it is likely that disruption of *Ddx3x* and *Eif2s3x* causes a fitness defect in female hepatocytes while male hepatocytes are functionally complemented by the Y chromosome paralogs. In this case, the sex-specific effect of these genes originates at a sex chromosome level. However, this approach is equally capable of identifying sex-specific effects that arise from hormonal or other differences between the sexes. Although uncovering sex-specific regulation of hepatocyte fitness was not the primary purpose of this study, these preliminary findings highlight the capacity to uncover such regulation by screening in the organism.

### Class I MHC and heparan sulfate biosynthesis are uniquely required for hepatocyte fitness in the organism

Perhaps the most valuable feature of genome-scale screening in the organism is the ability to investigate a phenotype in its native context. This captures all of the ways in which the cell interacts with the extracellular environment, many of which cannot be recapitulated in cell culture. To understand how hepatocyte fitness is regulated in the living organism, we first looked for patterns in the enriched and depleted genes by performing gene set enrichment analysis on the unified gene scores across the four mice. We did not identify any gene sets to be significantly enriched in our screen. However, we identified several gene sets that were significantly depleted in our screen (Figure 5A; Table S4). The gene sets included those previously established as essential for fitness in cell culture, including ribosome, proteasome, spliceosome, and RNA polymerase^1, 2, 17, 22, 27, 28^. However, we also identified several other gene sets not documented to be essential for cells in culture, including N-glycan biosynthesis, antigen processing/presentation, and glycosaminoglycan biosynthesis/heparan sulfate. Notably, these pathways all play major roles in the presentation or secretion of proteins at the cell surface^33, 34^ pointing to possible regulation of fitness from the extracellular environment that could only be identified by screening in the organismal context.

**Figure 5.**
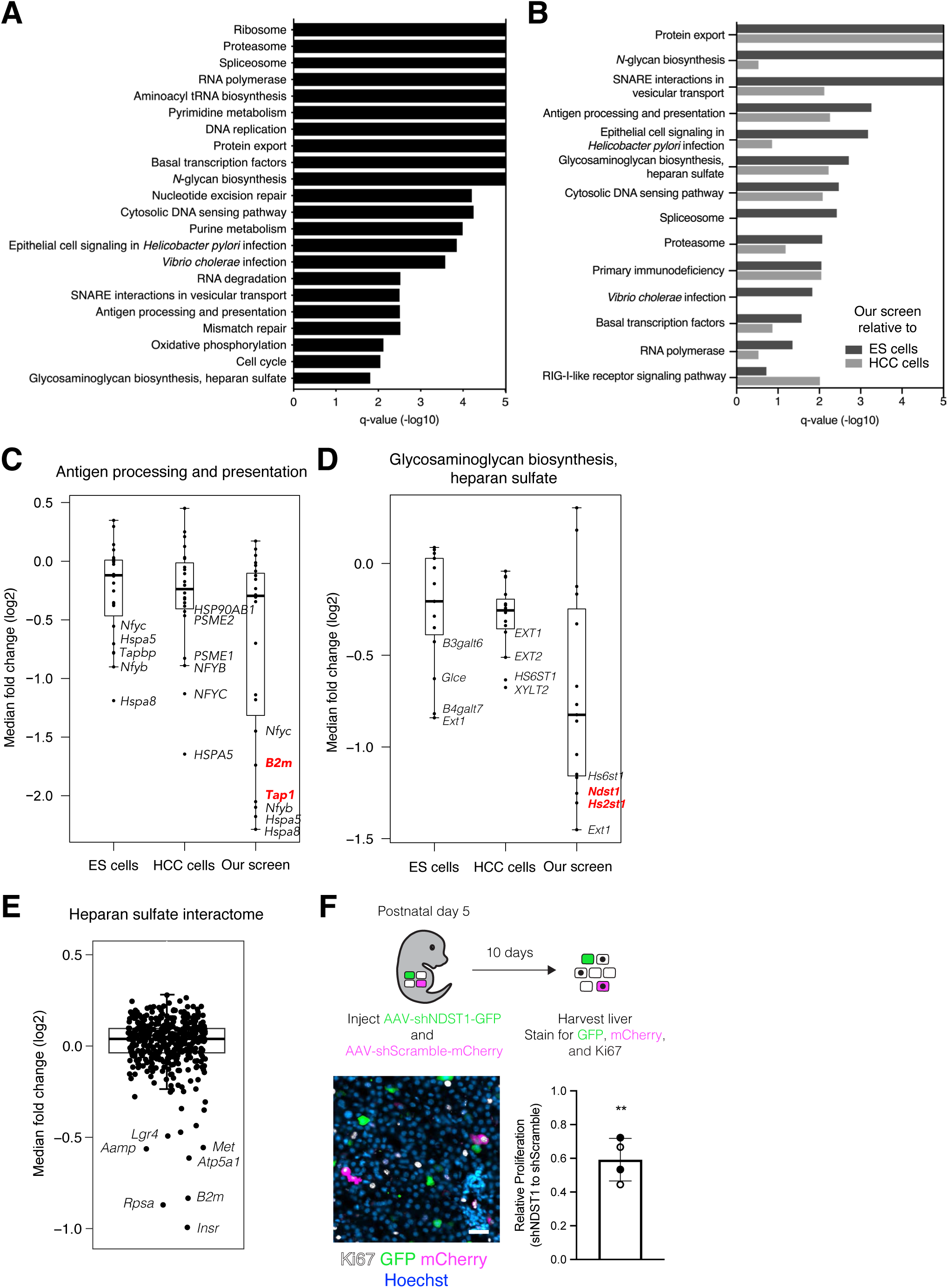
Class I MHC and heparan sulfate biosynthesis are uniquely required for hepatocyte fitness in the organism. (A) KEGG gene sets exhibiting significant depletion (FDR q-value < 0.05) at the endpoint of the screen ranked by FDR q-value (-log10). Bars extending to the end of the plot indicate an FDR q-value of 0. (B) KEGG gene sets exhibiting significant depletion (FDR q-value < 0.05) in our screen relative to screens in either mouse embryonic stem (ES) cells (dark grey bars) or human hepatocellular carcinoma (HCC) cell lines (light grey bars) ranked by FDR q-value (-log10). Bars extending to the end of the plot indicate an FDR q-value of 0. (C) Quantile-normalized median fold change (log2) for genes in the KEGG gene set for antigen processing and presentation in ES cell screens, HCC cell line screens, and our screen. Genes uniquely depleted in our screen are highlighted in red. The bounds of the box indicate the first and third quartiles, and the whiskers extend to the furthest data point that is within 1.5 times the interquartile range. (D) Quantile-normalized median fold change (log2) for genes in the KEGG gene set for glycosaminoglycan biosynthesis and heparan sulfate in ES cell screens, HCC cell line screens, and our screen. Genes uniquely depleted in our screen are highlighted in red. The bounds of the box indicate the first and third quartiles, and the whiskers extend to the furthest data point that is within 1.5 times the interquartile range. (E) Median fold change (log2) for genes in the heparan sulfate interactome in our screen. The bounds of the box indicate the first and third quartiles, and the whiskers extend to the furthest data point that is within 1.5 times the interquartile range. (F) Scheme for determining effects of NDST1 knockdown on hepatocyte proliferation (top panel). Image of liver from postnatal day 15 mouse injected with 1 x 10^10^ GC of AAV-shNDST1 and AAV-shScramble on postnatal day 5 immunostained for Ki67 (white), GFP (green) and mCherry (magenta) and counterstained for Hoechst (blue) (left panel). Scale bar, 25 µm. Quantification of proliferation as inferred by Ki67 positivity in shNDST1 hepatocytes relative to shScramble hepatocytes (right panel). Bar and whiskers indicate mean and standard deviation across mice, respectively, and closed and open circles represent values from male and female mice, respectively. n = 2 male and 2 female mice and 200 cells per shRNA per mouse. **p = 0.0023 by two-tailed Fisher’s exact test. See also Figure S5.

To determine whether any genes and pathways were indeed uniquely required for cell fitness in the organismal context, we compared our screen to screens of mouse embryonic stem (ES) cells in culture^25, 26^ and human hepatocellular carcinoma (HCC) cell lines in culture^27, 28^. We specifically chose ES cells and HCC cell lines in an effort to control for any species-specific and cell lineage-specific requirements for cell fitness. We identified four gene sets that were preferentially depleted in our screen relative to the cell culture screens: protein export, SNARE interactions in vesicular transport, antigen processing/presentation, and glycosaminoglycan biosynthesis/heparan sulfate (Figure 5B; Table S4). For the protein export and SNARE interactions gene sets, the most depleted genes in each set were depleted across all screens but to a greater extent in our screen (Figures S5A and S5B). However, for the antigen processing/presentation and glycosaminoglycan biosynthesis/heparan sulfate gene sets, some genes were depleted exclusively in our screen, suggesting a unique requirement for aspects of these pathways in hepatocyte fitness in the organism (Figures 5C and 5D).

Within the antigen processing/presentation gene set, *Tap1* and *B2m* were uniquely and dramatically depleted in our screen, ranking as the 41^st^ and 320^th^ most depleted genes, respectively (Figure 5C). Both of these genes are involved in—and required for—presentation of antigens at the cell surface by the class I major histocompatibility complex (MHC) pathway^34^. Indeed, within the antigen processing/presentation gene set, eight of the 32 genes attributed to the class I MHC pathway were depleted in our screen whereas none of the 13 genes attributed to the class II MHC pathway exhibited depletion (FDR < 0.25, Figure S5C). The class I MHC pathway presents intracellular antigens at the cell surface. At the cell surface, class I MHC can interact with both cytotoxic CD8 T cells and natural killer (NK) cells. The latter interaction can provide a pro-survival role by preventing NK cell cytotoxicity. However, loss of class I MHC alone should not be sufficient to induce NK cell cytotoxicity. Classically, NK cell activation requires both a loss of inhibitory signals, via loss of class I MHC, and presence of activating signals, expressed on the surface of transformed or infected cells^35^. Although our screening approach involves viral infection of hepatocytes, any inflammation resulting from lentiviral and AAV vectors has been shown to resolve within 72 hours^36, 37^ and we indeed did not observe any liver inflammation in our system (Figures S2E-G). This result suggests that class I MHC may play an essential survival role even in untransformed and uninflamed cells in the organism.

Within the glycosaminoglycan biosynthesis/heparan sulfate gene set, *Hs2st1* and *Ndst1* were uniquely depleted in our screen (Figure 5D). These two genes encode enzymes involved in the biosynthesis of heparan sulfate, a glycosaminoglycan conjugated to plasma membrane or extracellular matrix proteins to form heparan sulfate proteoglycans (HSPGs)^38^. HSPGs can be classified into three groups based on their location: cell membrane (syndecans, glypicans), extracellular matrix (agrin, perlecan, type XVIII collagen), and secretory vesicles (serglycin). To determine whether the requirement for heparan sulfate reflected the requirement for a specific HSPG, we analyzed the performance of individual HSPGs in our screen. No single HPSG was essential, suggesting that the requirement for heparan sulfate arises from a redundant function performed by multiple HSPGs (Figure S5D). Indeed, the syndecans are known to have redundant function. These transmembrane HSPGs play a variety of roles including facilitating attachment of cells to the extracellular matrix, protecting cytokines and growth factors from proteolysis, and serving as co-receptors for other transmembrane receptors. As such, these HSPGs interact with hundreds of other proteins in the extracellular matrix and at the cell surface. To gain insight into whether these interactions may be essential for hepatocyte fitness, we analyzed how proteins established to interact with HSPGs performed in our screen^39^. We observed a handful of heparan sulfate interacting genes for which knockout had significant fitness defects in hepatocytes (Figure 5E). Three of these, *Insr*, *Met*, and *Lgr4*, encode receptors for growth factors known to stimulate hepatocyte proliferation, suggesting that the requirement for HSPGs might be via cell-autonomous enhancement of growth factor signaling.

To examine the consequences of disrupted heparan sulfate biosynthesis by an alternative method, we used an orthologous AAV vector system in which we could deliver shRNA targeting NDST1 and thereby knockdown NDST1 protein in the whole liver or individual hepatocytes (Figure S5E). To determine whether loss of Ndst1 impaired hepatocyte proliferation, we injected neonatal mice with a low-dose mixture of AAV encoding shRNA targeting NDST1 alongside a GFP reporter (AAV-shNDST1) and AAV encoding a scramble shRNA alongside an mCherry reporter (AAV-shScramble) such that rare hepatocytes were expressing one of the shRNAs. We compared the proliferation of cells expressing shRNA targeting NDST1 to those expressing scramble shRNA within the same liver by immunostaining for the proliferation marker Ki67 alongside the fluorescent reporters. We observed a significant reduction in proliferation in the NDST1 knockdown hepatocytes compared to control knockdown hepatocytes in agreement with our screen results (Figure 5F). We therefore find that an enzyme in the heparan sulfate biosynthesis pathway is required for hepatocyte proliferation in the liver, perhaps via the role of HSPGs in potentiating mitogenic signaling in hepatocytes.

## DISCUSSION

Herein, we successfully developed, to our knowledge, the first method for genome-wide CRISPR screening directly within a single mouse. Our approach involves lentiviral delivery of a genome-wide sgRNA library to a neonatal inducible Cas9 mouse followed by Cas9 induction and phenotypic selection at any point in the animal’s lifetime. We applied this approach to uncover regulators of hepatocyte fitness in the neonatal liver. Our screen reliably identified positive and negative regulation of fitness in a single mouse with high reproducibility across mice. Not surprisingly, we found that hepatocytes in the neonatal liver share many of requirements for fitness with cells in culture. However, we also uncovered genes with sex-specific effects on hepatocyte fitness and genes that are uniquely required for hepatocyte fitness in the organism. Specifically, we found that hepatocytes in the liver, but not cells in culture, are dependent on the class I MHC and heparan sulfate biosynthesis pathways. Importantly, neither the sex-specific requirements nor the liver-specific requirements would have been discovered by screening a single cell line in culture. Our screen’s ability to uncover genetic regulation not identified in cell culture emphasizes the necessity and power of genome-wide screening in the living organism.

Our approach provides an adaptable and accessible method for the unbiased and comprehensive genetic dissection of diverse phenotypes within a living mouse. The stability of the sgRNA library in the liver and inducible nature of Cas9 allow for phenotypic selection at any point in the animal’s lifetime. This selection can be performed, as in our screen, by evaluating changes in the bulk hepatocyte population over time. Alternatively, selection could be performed by isolating hepatocytes from the liver and enriching for a single-cell phenotype. This combined flexibility of Cas9 induction and phenotypic selection makes this method a powerful tool for screening myriad processes spanning universal cellular phenomena, development and aging, hepatocyte-specific functions, and liver disease. Moreover, given the extent to which biological sex impacts physiology and disease, this ability to investigate the interaction between biological sex and gene function lends even further value to this technology. Importantly, these diverse applications are all within reach, as this versatile approach can be readily scaled, yet minimally requires a single mouse and fewer reagents than a cell culture screen.

Genome-scale CRISPR-Cas9 screening in the liver alone has the power to offer novel insights into diverse aspects of mammalian physiology and disease, but the full potential of high-throughput functional genomics in the organism lies in expanding this technology to other organs and CRISPR applications. Introducing this technology into other tissues will similarly be predicated on the ability to achieve stable, high-coverage sgRNA delivery in these tissues. This will require developing methods for efficient lentiviral delivery to organs beyond the liver and achieving genome-scale coverage in organs with much fewer cells. Once established in any organ, our overall approach can be readily adapted to incorporate other CRISPR-based techniques including CRISPR interference and activation. Our system therefore establishes the feasibility and foundation for genome-wide screening in a living organism. Building and expanding this platform will bring the experimental tractability once restricted to cell culture to the living organism, enabling unprecedented insight into mammalian physiology and disease.

## ACKNOWLEDGMENTS

We thank Mehreen Khan and Keya Viswanathan of the Whitehead Institute Functional Genomics Platform for technical assistance, Amanda Chilaka and Sumeet Gupta of the Whitehead Institute Genome Technology Core for high-throughput sequencing, Inma Barrasa of the Whitehead Institute Bioinformatics and Research Computing Core for RNA sequencing analysis, the Whitehead Institute Flow Cytometry Core, the Whitehead Institute Keck Microscopy Facility, the Koch Institute Swanson Biotechnology Center Microscopy Facility, and the Koch Institute Swanson Biotechnology Center Histology Facility. We thank Agilent Technologies for their commitment to supporting academic research. We thank Iain Cheeseman, Jonathan Weissman, Peter Reddien, Raghu Chivukula, Sharon Grossman, Tobiloba Oni, and members of the Knouse laboratory for their comments on the manuscript. This work was supported by the National Institutes of Health NIH Director’s Early Independence Award (DP5-OD026369) to K.A.K. and the National Cancer Institute Koch Institute Support Grant (P30-CA14051). K.A.K. was also supported by the Scott Cook and Signe Ostby Fund.

## AUTHOR CONTRIBUTIONS

H.R.K. and K.A.K. conceived the project, performed all experiments, analyzed the data, and wrote the manuscript.

## STAR METHODS

### RESOURCE AVAILABILITY

#### Lead Contact

Further information and requests for resources and reagents should be directed to and will be fulfilled by the Lead Contact, Kristin A. Knouse

#### Materials Availability

The plasmids and plasmid library generated in this study are in the process of being deposited at Addgene.

#### Data and Code Availability

The sequencing data generated in this study are in the process of being deposited at the GEO database.

### EXPERIMENTAL MODEL AND SUBJECT DETAILS

#### Animals

C57BL/6J mice (strain 000664) and LSL-Cas9 mice (strain 026175) were purchased from the Jackson Laboratory. Mice were either singly- or group-housed with a 12-hour light-dark cycle (light from 7 AM to 7 PM, dark from 7 PM to 7 AM) in a specific-pathogen-free animal facility with unlimited access to food and water. To deliver lentivirus, up to 100 µL of lentivirus in PBS was injected into the temporal vein of postnatal day one mice. For protein depletion tests, mice were injected with 1.25 x 10^7^ transduction units (TU) of sgRNA-mCherry lentivirus. For the screen, mice were injected with 5 x 10^7^ TU of sgRNA-mCherry lentiviral library. To deliver AAV-Cre, a stock solution of AAV8-TBG-Cre (Addgene 107787-AAV8) was diluted in PBS to a total volume of 20 µL and injected intraperitoneally into postnatal day five mice. For protein depletion tests and the screen, mice were injected with 2 x 10^11^ GC of AAV-TBG-Cre. To deliver AAV-shRNA, a stock solution of AAV8-mCherry-U6-scrmb-shRNA (AAV-shScramble, Vector Biolabs) and/or AAV8-GFP-U6-mNDST1-shRNA (AAV-shNDST1, Vector Biolabs) was diluted in PBS to a total volume of 20 µL and injected intraperitoneally into postnatal day five mice. To infect the entire liver to test protein depletion, mice were injected with 4 x 10^11^ GC of either AAV-shNDST1 or AAV-shScramble. To infect a subset of hepatocytes to compare proliferation, mice were injected with 1 x 10^10^ GC of both AAV-NDST1 and AAV-shScramble. All animal procedures were approved by the Massachusetts Institute of Technology Committee on Animal Care.

#### Cell lines

The mouse hepatocyte cell line AML12 was purchased from the American Type Culture Collection (ATCC) and cultured in DMEM/F12 medium supplemented with 10% fetal bovine serum, 10 µg/mL insulin, 5.5 µg/mL transferrin, 5 ng/mL selenium, and 40 ng/mL dexamethasone (ThermoFisher Scientific). HEK-293T cells were cultured in DMEM supplemented with 10% fetal bovine serum, 100 units/mL penicillin, and 100 µg/mL streptomycin (ThermoFisher Scientific). All cell lines were cultured at 37 °C with 5% CO_2_.

### METHOD DETAILS

#### Vector construction

The vector was produced through the following steps: 1) removal of the EFS-NS promoter and Cas9 from the parental vector and insertion of a hepatocyte-specific promoter driving dsRed expression, 2) replacement of dsRed with mCherry or mTurq2, and 3) removal of the puromycin resistance cassette.

To produce pLCv2-opti-stuffer-dsRed-puro, 100 ng of a synthetic gblock encoding the the HS-CRM8-TTRmin module^40^ upstream of dsRed (Integrated DNA Technologies) and 1 µg of LentiCRISPRv2-Opti^41^ (gift from David Sabatini, Addgene plasmid #163126), a lentiCRISPRv2 derivative containing an optimized scaffold (5’-GTTTAAGAGCTATGCTGGAAACAGCATAGCAAGTTT-3’)^42^ were digested sequentially with NheI and BamHI (New England Biolabs). The vector and fragment were purified using the QIAquick Gel Extraction Kit (Qiagen) and ligated with T4 DNA Ligase (New England Biolabs) in an 11 µL reaction to replace the EFS-NS promoter and Cas9 with the gblock fragment. 2.5 µL of the ligation was used to transform Stbl2 cells (Invitrogen) and DNA was isolated from ampicillin-resistant colonies with the QIAprep Spin Miniprep Kit (Qiagen). Clones were verified by Sanger sequencing (Quintara Biosciences) prior to retransformation and maxiprep using the ZymoPURE II Plasmid Maxiprep Kit (Zymo Research).

HS-CRM8-TTRmin-dsRed:

GAATTCGCTAGCACCGGCGCGCCGGGGGAGGCTGCTGGTGAATATTAACCAAGGTCACCCCAGTTATCGGAGGAGCAAACAGGGGCTAAGTCCACACGCGTGGTACCGTCTGTCTGCACATTTCGTAGAGCGAGTGTTCCGATACTCTAATCTCCCTAGGCAAGGTTCATATTTGTGTAGGTTACTTATTCTCCTTTTGTTGACTAAGTCAATAATCAGAATCAGCAGGTTTGGAGTCAGCTTGGCAGGGATCAGCAGCCTGGGTTGGAAGGAGGGGGTATAAAAGCCCCTTCACCAGGAGAAGCCGTCACACAGATCCACAAGCTCCTGACCGGTTCTAGAGCGCTGCCACCATGGTGCGCTCCTCCAAGAACGTCATCAAGGAGTTCATGCGCTTCAAGGTGCGCATGGAGGGCACCGTGAACGGCCACGAGTTCGAGATCGAGGGCGAGGGCGAGGGCCGCCCCTACGAGGGCCACAACACCGTGAAACTGAAGGTGACCAAGGGCGGCCCCCTGCCCTTCGCCTGGGACATCCTGTCCCCCCAGTTCCAGTACGGCTCCAAGGTGTACGTGAAGCACCCCGCCGACATCCCCGACTACAAGAAGCTGTCCTTCCCCGAGGGCTTCAAGTGGGAGCGCGTGATGAACTTCGAGGACGGCGGCGTGGTGACCGTGACCCAAGACTCCTCCCTGCAGGACGGCTGCTTCATCTACAAGGTGAAGTTCATCGGCGTGAACTTCCCCTCCGACGGCCCCGTAATGCAGAAGAAGACCATGGGCTGGGAGGCCTCCACCGAGCGCCTGTACCCCCGCGACGGCGTGCTGAAGGGCGAAATCCACAAGGCCCTGAAGCTGAAGGACGGCGGCCACTACCTGGTGGAGTTCAAGTCCATCTACATGGCCAAGAAGCCCGTGCAGCTGCCCGGCTACTACTACGTGGACTCCAAGCTGGACATCACCTCCCACAACGAGGACTACACCATCGTGGAGCAGTACGAGCGCACCGAAGGCCGCCACCACCTGTTCCTGGGATCCGGCG CAACAAACTTCTCTCTGCTGAAACAAGCCGGAGATGTCGAAGAGAATCCTGGACCGACCGAG

To construct pLCv2-opti-stuffer-mCherry-puro and pLCv2-opti-stuffer-mTurq2-puro, mCherry and mTurq2 were amplified from pKL028 and mTurquoise2-CMV (gifts from Iain Cheeseman), respectively, for 25 cycles with Q5 HotStart Polymerase (New England Biolabs) using the following primers (underlined nucleotides are homologous to the vector):

pLC_EBFP2_F: <u>GGTTCTAGAGCGCTGCCACC</u>ATGGTGAGCAAGGGCGAGGAG pLC_EBFP2_R: <u>GCCGGATCC</u>CTTGTACAGCTCGTCCATGCC

Amplicons and pLCv2-opti-stuffer-dsRed-puro were digested with XbaI and BamHI HF (New England Biolabs) and purified, ligated, transformed, and DNA was isolated and sequence verified as above.

To construct pLCv2-opti-stuffer-mCherry and pLCv2-opti-stuffer-mTurq2, a fragment encompassing the WPRE and 3’LTR was amplified from pLCv2-opti-stuffer-mCherry-puro as above using the following primers:

Puro_removal_F: TGAACGCGTTAAGTCGACAATCAACC Puro_removal_R: TCGAGGCTGATCAGCGGGTTTAAAC

The amplicon and pLCv2-opti-stuffer-mCherry-puro (or -mTurq2-puro) were digested with BsrGI-HF and PmeI (New England Biolabs) and purified as above. NEBuilder HiFi DNA Assembly Master Mix (New England Biolabs) was used to assemble 25 ng each of vector and fragment in a 20 µL reaction for 15 min at 50 °C. 50 µL of DH5-alpha cells were transformed with 2 µL assembly mix, and DNA was isolated and sequence verified as described above.

Individual sgRNAs were cloned as previously described^43^, using the following oligonucleotides (Integrated DNA Technologies):

sgAAVS1_F: CACCGGGGCCACTAGGGACAGGAT

sgAAVS1_R: AAACATCCTGTCCCTAGTGGCCCC

sgMaob_1_F: CACCGACGGATAAAGGATATACTTG

sgMaob_1_R: AAACCAAGTATATCCTTTATCCGTC

sgMaob_2_F: CACCGGGAAAATCATATGCCTTCAG

sgMaob_2_R: AAACCTGAAGGCATATGATTTTCCC

sgLmnb2_1_F: CACCGAGGTACGGGAGACCCGACGG

sgLmnb2_1_R: AAACCCGTCGGGTCTCCCGTACCTC

sgLmnb2_2_F: CACCGCTGCGCACCTACCTCACCGT

sgLmnb2_2_R: AAACACGGTGAGGTAGGTGCGCAGC

#### Lentivirus preparation, titration, and concentration

HEK-293T cells were seeded at a density of 750,000 cells/mL in 20 mL viral production medium (IMDM, Thermo Fisher Scientific, supplemented with 20% heat-inactivated fetal bovine serum, GeminiBio) in T175 flasks. After 24 hours, media was changed to fresh viral production medium. At 32 hours post-seeding, cells were transfected with a mix containing 76.8 µL Xtremegene-9 transfection reagent (Thermo Fisher Scientific), 3.62 µg pCMV-VSV-G (gift from Bob Weinberg, Addgene plasmid #8454), 8.28 µg psPAX2 (gift from Didier Trono, Addgene plasmid #12260), and 20 µg sgRNA plasmid in Opti-MEM (Thermo Fisher Scientific) to a final volume of 1 mL. Media was changed 16 hours later to 55 mL of fresh viral production medium. At 48 hours after transfection, virus was collected and filtered through a 0.45 µm filter, aliquoted, and stored at -80 °C until use.

To determine lentivirus titer, AML12 cells were transduced with a dilution series of lentivirus in the presence of 10 µg/mL polybrene for 16 hours. After four days, cells were harvested for flow cytometry analysis to determine percent of mTurq2- or mCherry-positive cells.

To concentrate lentivirus, lentiviral supernatant was ultracentrifuged at 23,000 RPM at 4°C for 2 hours in an SW 32 Ti swinging bucket rotor (Beckman Coulter). After centrifugation, media was decanted and pellets were air-dried at room temperature for 15 minutes. Pellets were then resuspended in PBS at room temperature for 30 minutes with gentle trituration. Concentrated lentivirus in PBS was stored for up to one week at 4°C prior to injection into mice.

#### Immunostaining

Livers were harvested and fixed in 4% paraformaldehyde in PBS at room temperature for 16-24 hours. Tissues were then washed with PBS and frozen in O.C.T. Compound (Tissue-Tek). Tissue sections of 12 to 30 µm thickness were prepared using a cryostat and adhered to Superfrost Plus Slides (Fisher Scientific). Slides were stored at -20 °C until use. To visualize endogenous mCherry and mTurq2 fluorescence, slides were dried at room temperature for 15 minutes, rehydrated in PBS for 5 minutes, permeabilized with 1% Triton X-100 in PBS for 15 minutes, and counterstained with Alexa Fluor 488 Phalloidin (ThermoFisher Scientific) diluted 1:500 in blocking buffer (3% bovine serum albumin and 0.3% Triton X-100 in PBS). To immunostain for endogenous proteins, slides were dried at room temperature for 4-24 hours and rehydrated in PBS for 5 minutes. Antigen retrieval was then performed by pressure cooking slides in sodium citrate buffer (10 mM tri-sodium citrate dihydrate, 0.05% Tween-20, pH 6.0) for 20 minutes in an Instant Pot (Amazon). Slides were rinsed in PBS for 5 minutes, dried briefly, and sections outlined with an ImmEdge hydrophobic pen (Vector Laboratories). Sections were permeabilized with 1% Triton X-100 in PBS for 15 minutes and blocked with blocking buffer for one hour. Sections were then incubated in primary antibodies diluted in blocking buffer at room temperature for 12-24 hours. Sections were washed with blocking buffer three times for 10 minutes each. Sections were then incubated in AlexaFluor secondary antibodies (ThermoFisher Scientific) diluted 1:1,000 in blocking buffer at room temperature for 1-2 hours. In some cases, 5 µg/mL Hoechst 33342 (ThermoFisher Scientific) was added to the secondary antibody solution. Sections were washed with blocking buffer twice for 10 minutes each followed by one wash with PBS for 5 minutes. Slides were then mounted in ProLong Gold Antifade reagent (ThermoFisher Scientific).

The following primary antibodies were used: Cas9 (1:200, clone 7A9-3A3, Abcam ab191468), asialoglycoprotein receptor 1 (ASGR1) (1:500, clone 114, Sino Biological 50083-R114), mCherry (1:500, clone 16D7, ThermoFisher Scientific M11217), monoamine oxidase B (MAO-B) (1:1,000, Novus Biologicals NBP1-87493), lamin B2 (1:1,000, clone EPR9701(B), Abcam ab151735), actin (1:250, clone AC-74, Sigma Aldrich A2228), CD45 (1:500, Abcam ab10558), Ki67 (1:200, clone SP6, Abcam ab16667), mCherry (1:2,000, Abcam ab167453), GFP (1:1,000, Abcam ab13970), and Ki67 (1:200, clone SolA15, ThermoFisher Scientific 14-5698-80). The Cas9 antibody was directly conjugated to AlexaFluor 647 using the AlexaFluor 647 Antibody Labeling Kit (ThermoFisher Scientific). The actin antibody was directly conjugated to DyLight 405 using the DyLight 405 antibody labeling kit (ThermoFisher scientific).

#### Image analysis

Images were acquired using either a CSU-22 spinning disc confocal head (Yokogawa) with Borealis modification (Andor) mounted on an Axiovert 200M microscope (Zeiss) with 10X or 40X objectives (Zeiss), an Orca-ER CCD camera (Hamamatsu), and MetaMorph acquisition software (Molecular Devices) or a McBain-Yokogawa spinning disk confocal head mounted on a Nikon Ti microscope with 20X objective (Nikon), a Clara CCD camera (Andor), and NIS Elements acquisition software (Nikon).

Images were analyzed using Volocity (Quorum Technologies). To determine the number of hepatocytes in the postnatal day one liver, liver volume was measured by volume displacement and hepatocyte volume was measured by immunostaining postnatal day one liver sections for the hepatocyte marker ASGR1 and actin. The proportion of a section occupied by ASGR1-positive cells was calculated to determine the proportion of liver volume comprised by hepatocytes and the hepatocyte volume was determined by measuring the x, y, and z dimensions of single hepatocytes, multiplying these three dimensions to calculate the volume of each hepatocyte, and averaging this volume across at least 29 cells per liver. To measure MAO-B and lamin B2 intensity, a single Z plane at the center of the cell was identified and the cytoplasm or nucleus was outlined to measure the signal intensity per µm. A similar procedure was done on sections stained only with secondary antibodies to calculate the average background intensity. This average background intensity was subtracted from each MAO-B and lamin B2 intensity measurement and the background-subtracted measurements were then normalized within a given sample (mCherry-positive or – negative hepatocytes within a single liver). To measure proliferation in AAV-shRNA-infected livers, GFP-positive and mCherry-positive hepatocytes were first identified on the basis of GFP and mCherry signal alone. Once identified, these hepatocytes were analyzed for their Ki67 signal and scored as positive or negative for Ki67.

#### Hematoxylin and eosin staining

Livers were harvested and fixed in 4% paraformaldehyde in PBS at room temperature for 16-24 hours. Livers were embedded in Paraplast X-tra paraffin (Leica Biosystems).

Tissue sections of 4 µm thickness were prepared using a microtome. Sections were stained with hematoxylin (3 minutes, Leica Biosystems) and eosin (10 seconds, Leica Biosystems) on a Tissue-Tek Prisma automated slide stainer (Sakura) and coverslipped on a Tissue Tek Glas g2 automated coverslipper (Sakura).

#### Immunoblotting

To prepare protein lysates from liver tissue, 50 mg of liver was homogenized in 1 mL of RIPA buffer (50 mM Tris pH 8.0, 150 mM sodium chloride, 1% NP-40, 0.5% sodium deoxycholate, 0.1% sodium dodecyl sulfate) containing cOmplete protease inhibitors (Roche) using a Bio-Gen PRO200 handheld homogenizer (PRO Scientific).

Homogenate was centrifuged at 18,000 G at 4 °C for 20 minutes. Supernatant was combined with 5X sample buffer (250 mM Tris pH 6.8, 50% glycerol, 0.025% bromophenol blue, 5% sodium dodecyl sulfate, 5% beta-mercaptoethanol). Samples were separated on homemade polyacrylamide gels and transferred to Immobilon-FL membranes (Millipore) via wet transfer. Membranes were blocked in 5% milk in TBST (50 mM Tris pH 8.0, 150 mM NaCl, 0.1% Tween-20) for 1 hour at room temperature. Membranes were incubated in primary antibody diluted in blocking solution at 4 °C with rocking overnight and washed with TBST for five minutes five times. Membranes were incubated in HRP-conjugated secondary antibody diluted in blocking solution at room temperature with rocking for one hour and washed with TBST for five minutes five times. Membranes were incubated in ECL Prime Western Blotting Detection Reagent (GE Healthcare) for five minutes and imaged on an ImageQuant LAS 4000 imager (GE Healthcare).

The following primary antibodies were used: Ndst1 (1:1,000, Sigma SAB1307040) and beta-actin (1:10,000, clone AC-74, Sigma A2228).

The following secondary antibodies were used: Rabbit (1:50,000, Abcam ab205718) and mouse (1:10,000, Abcam ab205719).

#### RNA sequencing

For surgical resection time points, partial hepatectomies were performed on 8 week-old mice as previously described^44^. For toxic injury time points, 8 week-old mice were injected intraperitoneally with 2 µL/gram of 25% carbon tetrachloride diluted in corn oil (Sigma Aldrich). For all time points, livers from three male C57BL/6J mice were harvested, flushed with PBS, immediately immersed in RNAlater (Qiagen), incubated at room temperature for 24 hours, and stored at -20 °C until future use. To isolate RNA, 30 mg of each tissue was removed from RNAlater and homogenized in 700 µL of QIAzol lysis reagent (Qiagen) using the TissueRuptor homogenizer (Qiagen). RNA was purified using the miRNeasy Kit (Qiagen) according to kit instructions and eluted in 30 µL of nuclease-free water. RNA sequencing libraries were prepared using KAPA mRNA HyperPrep Kit (KAPA Biosystems) according to manufacturer instructions. Briefly, 0.1-1 ug of total RNA was enriched for polyadenylated sequences using oligo-dT magnetic bead capture. The enriched mRNA fraction was then fragmented and first-strand cDNA generated using random primers. Strand specificity was achieved during second-strand cDNA synthesis by replacing dTTP with dUTP to quench the second strand during amplification. The resulting cDNA was A-tailed and ligated with indexed adapters. The library was amplified using a DNA polymerase that cannot incorporate past dUTPs to quench the second strand during PCR. The libraries were quantified using a KAPA qPCR Library Quantification Kit (KAPA Biosystems) as per manufacturer instructions. The samples were sequenced on a HiSeq 2500 (Illumina) based on qPCR concentrations. Base calls were performed by the instrument control software and further processed using the Offline Base Caller version 1.9.4 (Illumina). Samples were mapped with STAR version 2.6.1a^45^ to the mouse genome release mm10, using a gtf file from ENSEMBL version GRCm38.91, and setting the maximum intron length (“alignIntronMax”) parameter to 50000. We ran featureCounts version 1.6^46^ to assign reads to genes using the same gft file and setting “-s” parameter to 2. We normalized gene counts with DESeq2 version 1.22.2^47^. FPKMs were calculated using the function fpkm within the DESeq2 package. The FPKMs values for the three replicates were averaged, and protein coding genes were selected based on the annotation in the gtf file.

#### sgRNA library preparation

Genes with an average FPKM > 0.3 in any of the RNA sequencing time points were chosen to build a liver transcriptome-wide library. sgRNA sequences were designed using the Broad Institute GPP sgRNA Designer^18, 48^ using the Azimuth 2.0 rule set. For genes which were not identified by the program, alternative gene names from ENSEMBL versions GRCm38.76 - .93 were attempted. A small number of designed sgRNAs targeted multiple genes; the sgRNA names and gene names were manually annotated to indicate all targeted genes for these cases. Non-targeting and control-gene-targeting sgRNAs from Doench et al. 2014^18^ were also included. sgRNA sequences from this control set that were identical to a sequence already in our library were annotated according to the targeted gene; those that did not overlap with sequences in our sgRNA library were annotated as control sgRNAs. The library contains 71,878 sgRNAs targeting 13,189 genes.

For sgRNAs beginning with a nucleotide other than G, a G was prepended. The following adapters were added to all sgRNA sequences:

Upstream: TATCTTGTGGAAAGGACGAAACACC Downstream: GTTTAAGAGCTATGCTGGAAACAGCATAGC

Multiple rounds of cloning were combined to generate the final plasmid library. The oligonucleotide library (Agilent Technologies) was amplified for 16 cycles using Q5 HotStart Polymerase (New England Biolabs) using a gradient annealing temperature ranging from 50-62 °C across 8, 50 µL reactions using the forward primer TTTCTTGGCTTTATATATCTTGTGGAAAGGACGAAACACC and the reverse primer ATTTAAACTTGCTATGCTGTTTCCAGCATAGCTCTTAAAC and the following program:

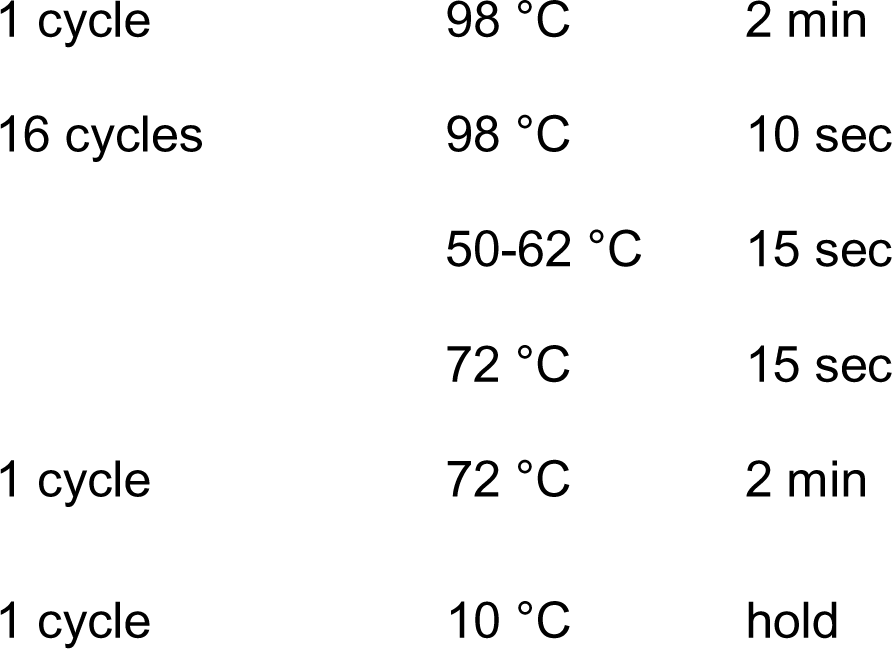

Reactions were pooled and purified by DNA Clean and Concentrator 5 (Zymo Research). pLCv2-opti-stuffer-mCherry was digested as described^43^, and either gel purified using a Zymoclean Gel DNA Recovery Kit (Zymo Research) followed by Ampure XP bead purification (Beckman Coulter) or DNA Clean and Concentrator 5. The library was assembled using NEBuilder HiFi DNA Assembly Master Mix (New England Biolabs) in 4 x 20 µL reactions at 50 °C for 1 hour using 100 ng of vector per 5-10 ng of PCR amplicon. The reactions were combined and 2.5 µL of the assembly reaction or a control reaction without amplicon were used to transform NEB5-alpha cells (New England Biolabs) to measure background assembly. Subsequently, the assembly reactions were combined, concentrated using Ampure XP beads, resuspended in 8 µL water, and used to electroporate 1-4 tubes of Endura DUO electrocompetent cells (Lucigen) at 1.8 kV distributed over 2 cuvettes (0.1 cm gap width) per tube using a Micropulser Electroporator (Bio-Rad Laboratories). 10-fold serial dilutions of a 10 µL aliquot were plated on LB plates with ampicillin at 100 µg/mL to assess electroporation efficiency, and the remainder of each electroporation (2 cuvettes) was plated on LB agar supplemented with 100 µg/mL ampicillin in 4 x 245 mm square bioassay dishes (Corning). Plates were incubated overnight at 30 °C and colonies were scraped the next morning. DNA was isolated using the ZymoPURE II Plasmid Maxiprep Kit (Zymo Research). Plasmid DNA from multiple rounds of assembly and electroporation were combined according to the measured electroporation efficiency to achieve 25-fold coverage of the library. sgRNA representation was measured by high-throughput sequencing as described below.

To improve coverage of some of the sgRNAs in the library, a second library containing ∼7,500 sgRNAs was synthesized and cloned as above, with the following modifications: assembly was performed using NEB Gibson Assembly mix (New England Biolabs) using a ratio of 200 ng vector : 10 ng sgRNA in each 20 µL reaction, and the final combined and concentrated reaction was used to electroporate a single tube of Endura DUO cells.

Subsequent propagation of the plasmid library was performed using 50 ng plasmid library per single tube of Endura DUO cells.

All steps were performed according to manufacturer’s instructions, except where noted.

#### Genomic DNA isolation

Livers were harvested from mice, separated into individual lobes, minced into 15 mg pieces using a razor blade, snap-frozen in liquid nitrogen, and stored at -80 °C until use. Genomic DNA (gDNA) was isolated from livers using the illustra blood genomicPrep Mini Spin Kit (Cytiva) using one column for every 7.5 mg of tissue. The manufacturer’s protocol was used with the following modifications: 20 µL of 10 mg/mL Proteinase K (Millipore-Sigma) solution in water was added per 7.5 mg of tissue. Tissue was disrupted by thoroughly pipetting prior to adding lysis buffer, vortexing, and incubating at 56 °C overnight. Elution was performed using 25 µL of water pre-heated to 70 °C. Samples were combined by lobe and concentration was measured using the Qubit dsDNA HS Assay Kit (Invitrogen).

For the induction, equal amounts of gDNA from each lobe were combined within each mouse, and equal inputs from four mice were combined to prepare a single sequencing library. For the endpoint, gDNA from each lobe within a mouse was combined proportionally to the average lobe mass across mice measured at liver harvest. A sequencing library was prepared for each mouse individually using equal total gDNA input per mouse.

#### Sequencing library preparation and DNA sequencing

All PCR reactions were performed in 50 µL reactions using ExTaq Polymerase (Takara Bio) with the following program:

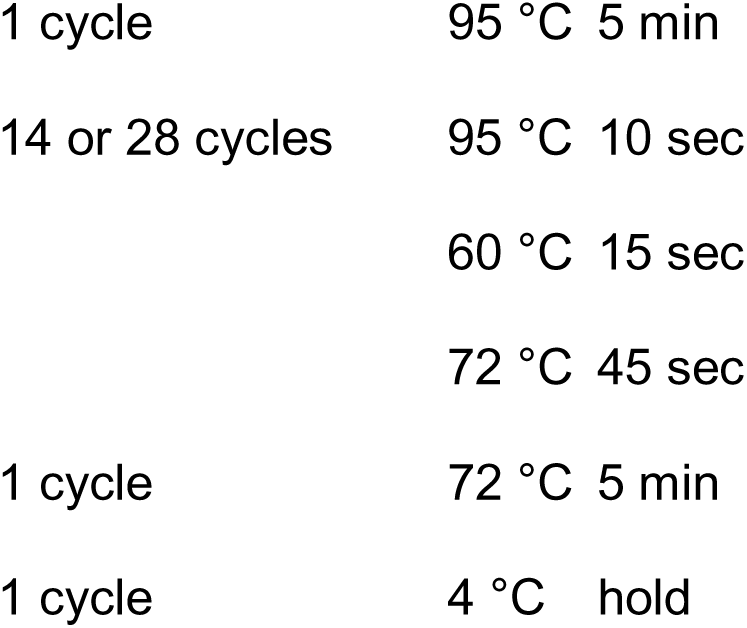

Using the following primers:

Forward: AATGATACGGCGACCACCGAGATCTACACCGACTCGGTGCCACTTTT

Reverse: CAAGCAGAAGACGGCATACGAGATCnnnnnnTTTCTTGGGTAGTTTGCAGTTTT

Where “nnnnnn” denotes the barcode used for multiplexing. 10 ng of plasmid DNA was amplified for 14 cycles in 4 x 50 µL reactions. 1, 3, or 6 µg of gDNA was initially amplified for 28 cycles in 50 µL test PCR reactions. Subsequently, 226 µg of gDNA (induction) was used in 38 reactions, or 75 µg of gDNA (endpoint) was used in 25 reactions per mouse. All reactions were cleaned and concentrated using Ampure XP beads prior to sequencing for 50 cycles on an Illumina Hiseq 2500 using the following primers:

Read 1 sequencing primer: GTTGATAACGGACTAGCCTTATTTAAACTTGCTATGCTGTTTCCAGCATAGCTCTTA AAC

Index sequencing primer: TTTCAAGTTACGGTAAGCATATGATAGTCCATTTTAAAACATAATTTTAAAACTGCAA ACTACCCAAGAAA

Base calls were performed by the instrument control software and further processed using the Offline Base Caller (Illumina) v. 1.9.4.

#### Screen analysis

For initial measurement of sgRNA representation in the plasmid library or induction time point, sequencing reads were mapped to the library, each sgRNA was given a pseudocount of 1, and reads per million (RPM) was calculated as previously described^17^. Raw counts were processed using MAGeCK for downstream analysis^19^. The plasmid library and induction timepoint were used as control samples and to estimate variance, and each endpoint mouse was processed separately. For mouse 4, sgLmnb2_1 was removed prior to MAGeCK analysis, as the high representation of this sgRNA (an sgRNA used for development of the screening method) was likely due to contamination during sequencing library preparation. Counts data from Tzelepis et al.^25^ (day 14) and Shohat et al.^26^ (day 18) were processed individually using MAGeCK. The corresponding plasmid libraries were used as control samples. For our screen, Shohat et al., and Tzelepis et al., the null distribution was generated using the control sgRNA set (Supplementary Table 2). For Tzelepis et al., the three replicate day 14 samples were processed together to generate a single gene score, and those three samples were used to estimate variance. For all screens, the gene test FDR threshold was set to 0.05, the sgRNA p-value was FDR-adjusted, and the gene score was calculated using the median. Twenty-four human hepatocellular carcinoma screens from the CRISPR (Avana) Public 20Q4 release were downloaded from the Broad DepMap portal using “Liver” as a lineage filter and “Hepatocellular Carcinoma” as a lineage subtype filter^27, 28^.

All downstream analyses were performed in R version 3.6.0 or 4.2.1, and all plots were generated in either base R, using the R corrplot package, or in GraphPad Prism Version 7.0d. For comparisons within our screen, the gene scores from individual mice were not normalized across mice, as each mouse serves as a replicate screen. The gene score for each gene across mice was tested against all gene scores using an unpaired two-sample Wilcoxon test. The p-values from this test were adjusted using the Benjamini-Hochberg (FDR) procedure. The median log2 fold change across mice was used as input for pre-ranked GSEA using the c2.cp.kegg.v7.1.symbols.gmt gene sets. For comparisons between screens from different sources, all screens from all the sources in the specific comparison were quantile normalized to one another using the preprocessCore R package prior to calculating the median log2 fold change within the screens from each source. This normalized median log2 fold change was subtracted from the normalized median log2 fold change of our screens to generate a differential score used as input for pre-ranked GSEA using the c2.cp.kegg.v7.1.symbols.gmt gene sets. For converting mouse gene symbols to human gene symbols, the Mouse_Gene_Symbol_Remapping_to_Human_Orthologs_MSigDB.v7.1.chip was used (note that this excludes genes that have multiple annotations in either human or mouse).

Pearson correlation was used to compare gene effects between mice. Spearman correlation was used to compare gene effects with liver mRNA expression and protein half-life in hepatocytes^16^. The TUSON dataset of predicted tumor suppressor genes was sorted by ascending FDR q-value and the top 50 genes present in the compared datasets were used^29^. Distribution differences were tested and p-values were calculated using the Kolmogorov-Smirnov test. A one-sided test was used for gene sets for which a phenotype could be predicted (core essential genes, tumor suppressor genes, and control enriched and depleted genes); all other comparisons used a two-sided test.

For comparison of sex-specific fitness effects, only genes with an average of > 2 sgRNAs detected across mice were considered. For sex-specific enriched genes, a median fold change (log2) > 0.5 across two mice of a given sex and an absolute median fold change (log2) difference of > 0.25 compared to the other sex was required. To identify true tumor-suppressor-like genes, a median fold change (log2) > -0.5 was required in mice of the other sex. For sex-specific depleted genes, a median fold change (log2) of < -0.5 across two mice of a given sex and an absolute median fold change (log2) difference of > 0.75 compared to the other sex was required. To identify true sex-specific essential genes, a median fold change (log2) > -0.5 was required in mice of the other sex.

### QUANTIFICATION AND STATISTICAL ANALYSIS

#### Statistical analysis

The statistical details for any given experiment are provided in the corresponding figure legend. Additional information about statistical analysis can be found in the relevant Method Details sections.

#### Software

STAR version 2.6.1a Conda version 4.9.2 MAGeCK-RRA version 0.5.9.2 R version 3.6.0 or 4.2.1

featureCounts version 1.6

DESeq2 version 1.22.2

preprocessCore version 1.48.0

corrplot version 0.84

GSEA version 4.1.0

Mouse_Gene_Symbol_Remapping_to_Human_Orthologs_MSigDB.v7.1.chip

Human_Symbol_with_Remapping_MsigDB.v7.1.chip c2.cp.kegg.v7.1.symbols.gmt

GraphPad Prism version 7.0d

## SUPPLEMENTAL INFORMATION

**Supplemental Figure 1.**
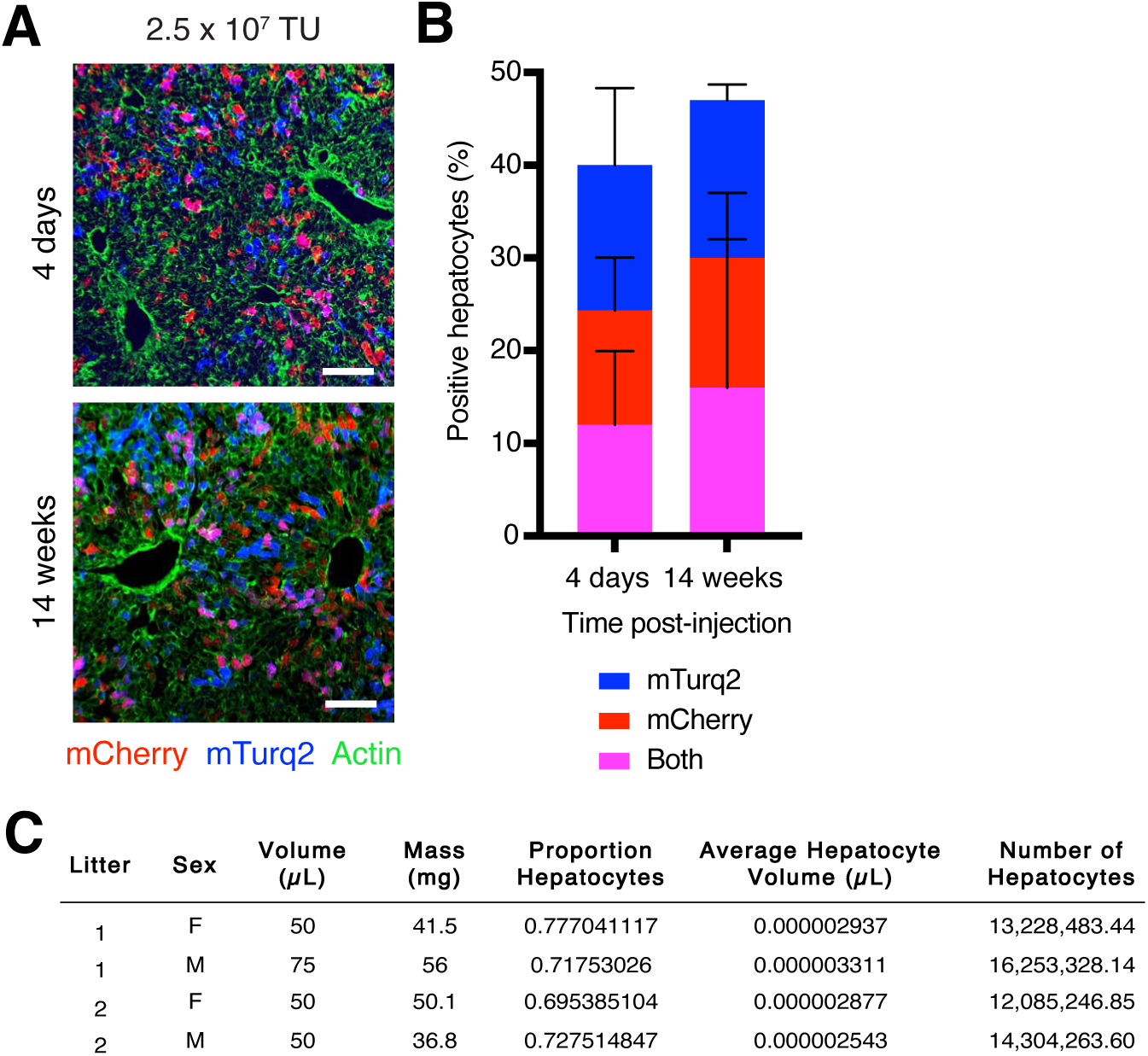
Genome-scale sgRNA delivery in a single mouse liver, Related to Figure 1. (A) Images of endogenous mCherry and mTurq2 fluorescence in livers from mice four days or 14 weeks after injection of 2.5 x 10^7^ TU of an equal mixture of sgAAVS1-mCherry and sgAAVS1-mTurq2 lentiviruses. Scale bars, 100 µm. (B) Percent mCherry-, mTurq2-, and double-positive hepatocytes in livers from mice four days or 14 weeks after injection of 2.5 x 10^7^ TU of an equal mixture of sgAAVS1-mCherry and sgAAVS1-mTurq2 lentiviruses. Error bars indicate standard deviation. n = 3 mice per time point and 200 hepatocytes per mouse. (C) Table of values used to estimate the number of hepatocytes in postnatal day one livers. Liver volume was measured by volume displacement and percent hepatocytes and hepatocyte volume were measured by immunostaining and microscopy.

**Supplemental Figure 2.**
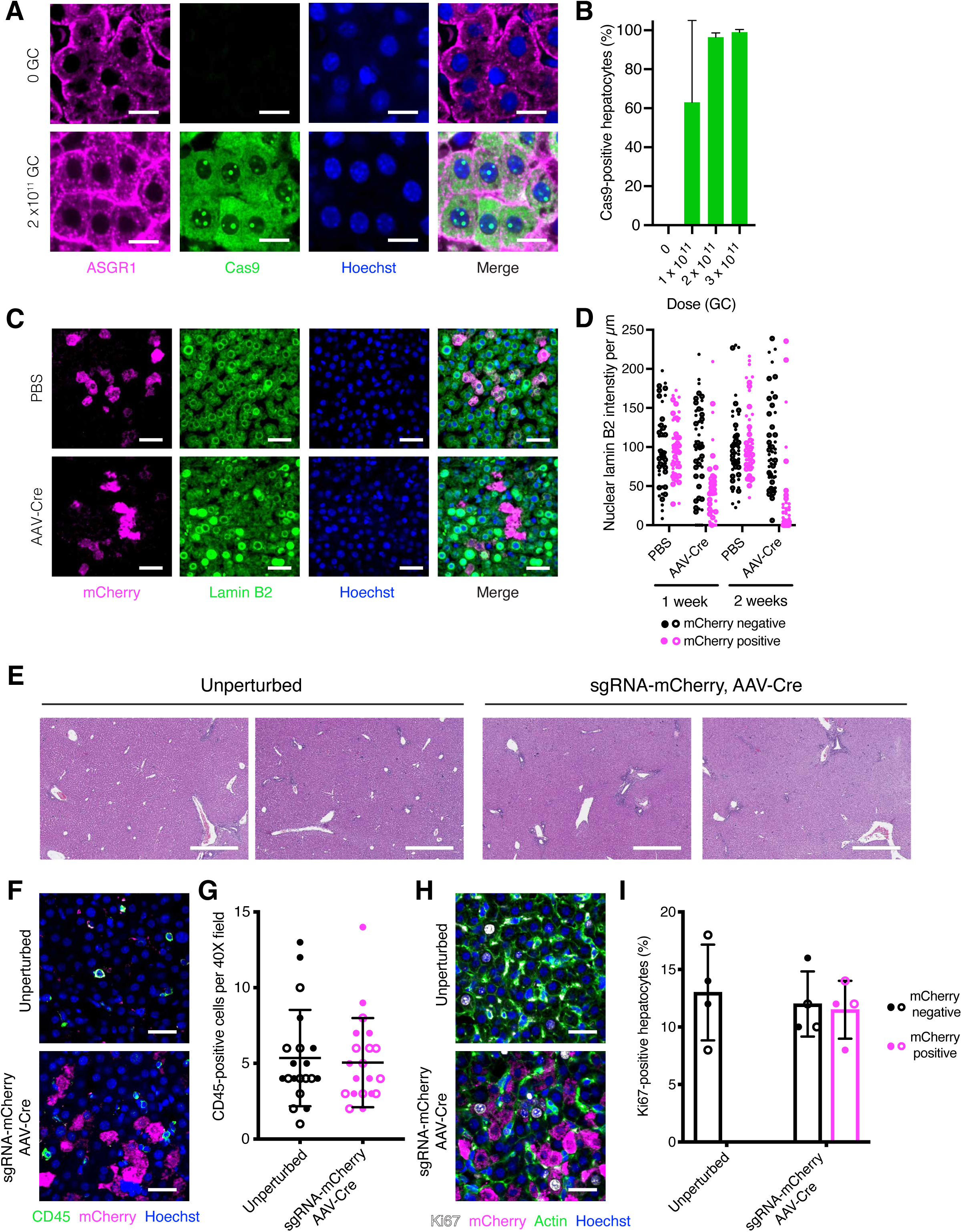
Temporally-controlled protein depletion in the mouse liver, Related to Figure 2. (A) Images of livers from LSL-Cas9 mice injected with 0 or 2 x 10^11^ GC of AAV-Cre on postnatal day five and harvested four days thereafter immunostained for ASGR1 (hepatocyte marker, magenta) and Cas9 (green) and counterstained with Hoechst (blue). Scale bars, 15 µm. (B) Percent Cas9-positive hepatocytes in livers from LSL-Cas9 mice injected with varying doses of AAV-Cre on postnatal day five and harvested four days thereafter as determined by Cas9 immunostaining. Error bars indicate standard deviation. n = 1 male and 1 female mouse per dose and 200 hepatocytes per mouse. (C) Images of livers from LSL-Cas9 mice injected with sgLmnb2-mCherry followed by PBS or AAV-Cre immunostained for mCherry (magenta) and lamin B2 (green) and counterstained with Hoechst (blue). Scale bars, 45 µm. (D) Nuclear lamin B2 intensity per µm in mCherry-positive and mCherry-negative hepatocytes from LSL-Cas9 mice injected with sgLmnb2-mCherry followed by PBS or AAV-Cre. Closed and open circles represent values from male and female mouse, respectively. n = 1 male and 1 female mouse per condition and 25 cells per mouse. (E) Images of livers from unperturbed postnatal day 12 LSL-Cas9 mice or postnatal day 12 LSL-Cas9 mice injected with 1.25 x 10^7^ TU of sgMaob-mCherry or sgLmnb2-mCherry lentivirus on postnatal day 1 followed by AAV-Cre on postnatal day 5 stained with hematoxylin and eosin. Left and right images represent one male and one female mouse, respectively, from each condition. Scale bars, 500 µm. (F) Images of livers from unperturbed postnatal day 12 LSL-Cas9 mice or postnatal day 12 LSL-Cas9 mice injected with 1.25 x 10^7^ TU of sgMaob-mCherry or sgLmnb2-mCherry lentivirus on postnatal day 1 followed by AAV-Cre on postnatal day 5 immunostained for CD45 (green) and mCherry (magenta) and counterstained for Hoechst (blue). Scale bars, 45 µm. (G) Number of CD45-positive cells per 40X field in unperturbed postnatal day 12 LSL-Cas9 mice or postnatal day 12 LSL-Cas9 mice injected with 1.25 x 10^7^ TU of sgMaob-mCherry or sgLmnb2-mCherry lentivirus on postnatal day 1 followed by AAV-Cre on postnatal day 5 as determined by CD45 immunostaining. Bar and whiskers indicate mean and standard deviation across mice, respectively, and closed and open circles represent values from male and female mice, respectively. n = 2 male and 2 female mice per condition and five fields per mouse. (H) Images of livers from unperturbed postnatal day 12 LSL-Cas9 mice or postnatal day 12 LSL-Cas9 mice injected with 1.25 x 10^7^ TU of sgMaob-mCherry or sgLmnb2-mCherry lentivirus on postnatal day 1 followed by AAV-Cre on postnatal day 5 immunostained for Ki67 (white), mCherry (magenta) and actin (green) and counterstained for Hoechst (blue). Scale bars, 45 µm. (I) Percent Ki67-positive hepatocytes in livers from unperturbed postnatal day 12 LSL-Cas9 mice or postnatal day 12 LSL-Cas9 mice injected with 1.25 x 10^7^ TU of sgMaob-mCherry or sgLmnb2-mCherry lentivirus on postnatal day 1 followed by AAV-Cre on postnatal day 5 as determined by Ki67 immunostaining. Bar and whiskers indicate mean and standard deviation across mice, respectively, and closed and open circles represent values from male and female mice, respectively. n = 2 male and 2 female mice per condition and 50 cells per mouse.

**Supplemental Figure 3.**
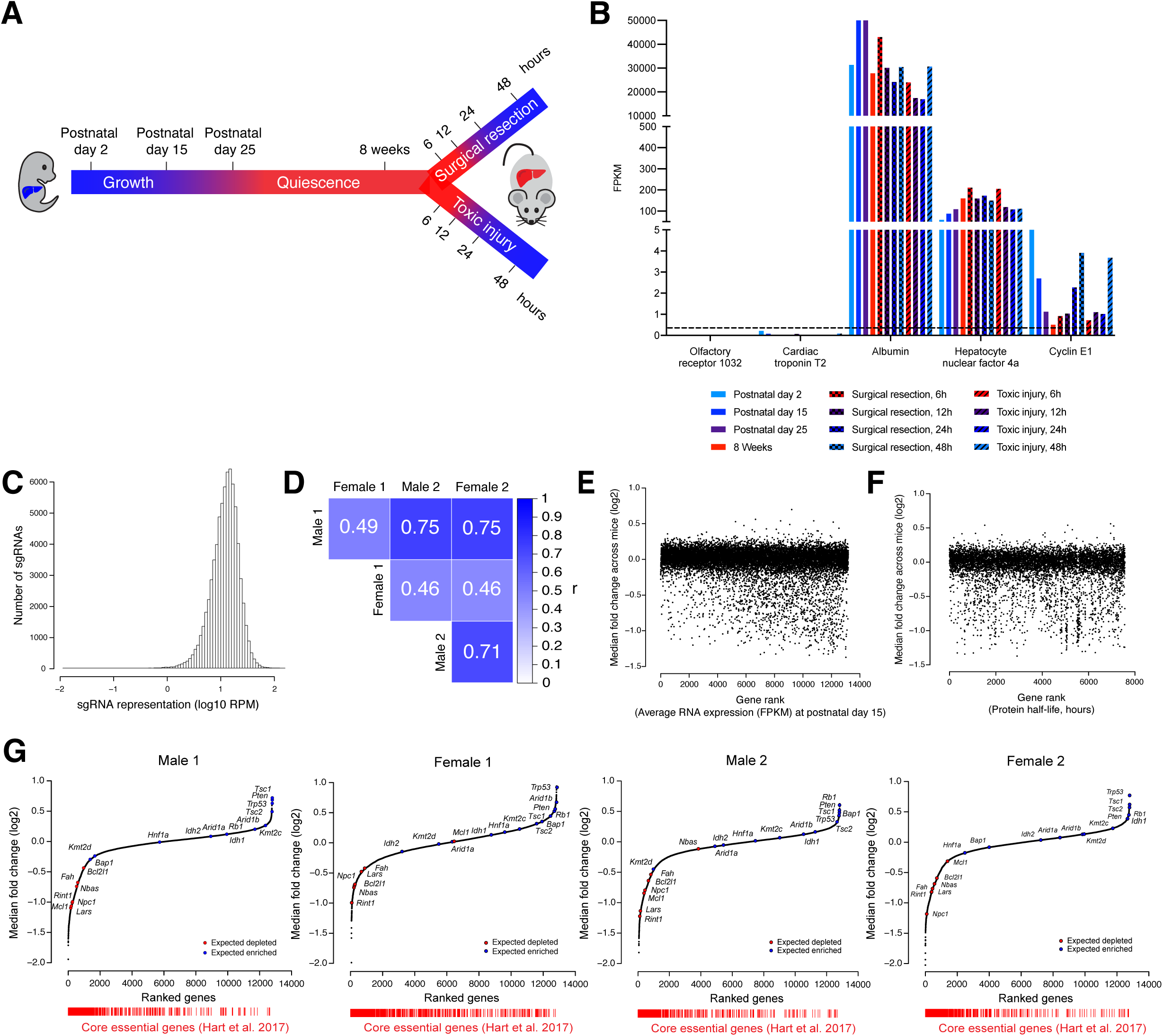
A genome-scale screen for hepatocyte fitness in the neonatal mouse liver, Related to Figure 3. (A) Time points during liver growth, quiescence, and regeneration from which livers were harvested for RNA sequencing. Partial hepatectomy and carbon tetrachloride were used as surgical resection and toxic injury models, respectively. (B) Average RNA expression (FPKM) of representative protein-coding genes at different time points of liver growth, quiescence, and regeneration. Dashed line indicates FPKM cutoff of 0.3. n = 3 male mice per time point. (C) Number of sgRNAs with a given representation (log10 RPM) for all sgRNAs in the library. (D) Pearson correlation (r) for each plot in Figure 3D. (E) Median fold change across mice (log2) of genes ranked by average RNA expression (FPKM) at postnatal day 15 for each gene. Two-sided Spearman ρ = -0.15, p < 2.2 x 10^-16^ (F) Median fold change across mice (log2) of genes ranked by protein half-life (hours) for each gene. Two-sided Spearman ρ = -0.05, p = 0.003. (G) Genes ranked by median fold change (log2) in each of the four mice. Highlighted are control gene sets consisting of tumor suppressor genes in hepatocellular carcinoma (expected to enrich, blue) and genes required for hepatocyte viability (expected to deplete, red). Core essential genes (red bars) are positioned below based on gene rank to demonstrate their significant depletion in each mouse. Expected gene depletion p = 1 x 10^-5^, 1.6 x 10^-4^, 1.4 x 10^-4^, and 1 x 10^-5^ for male 1, female 1, male 2, and female 2, respectively, by one-sided Kolmogorov-Smirnov test, expected gene enrichment p = 0.0032, 0.0052, 0.0053, and 0.0039 for male 1, female 1, male 2, and female 2, respectively, by one-sided Kolmogorov-Smirnov test. Core essential gene depletion p < 2.2 x 10^-16^ for each mouse by one-sided Kolmogorov-Smirnov test.

**Supplemental Figure 4.**
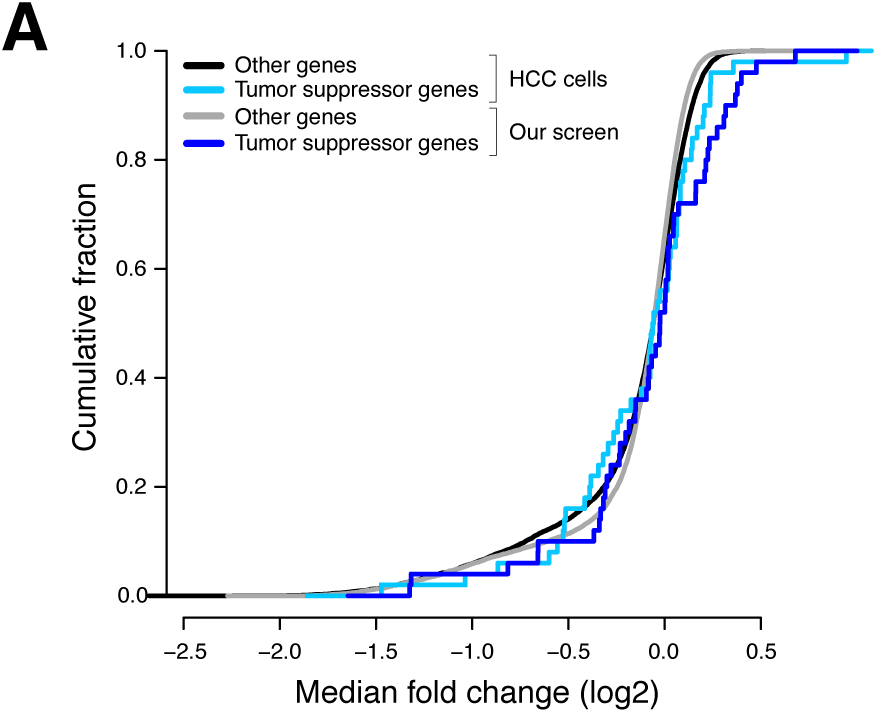
Genome-scale screening in the organism enhances discovery of tumor suppressor genes and uncovers genes with sex-specific effects, Related to Figure 4. (A) Cumulative fraction of tumor suppressor genes (cyan, blue) and other genes (black, grey) based on quantile-normalized median fold change (log2) of their gene scores across screens in human hepatocellular carcinoma (HCC) cell lines and our screen. HCC cells p > 0.05 by one-sided Kolmogorov-Smirnov test, our screen p = 0.0028 by one-sided Kolmogorov-Smirnov test.

**Supplemental Figure 5.**
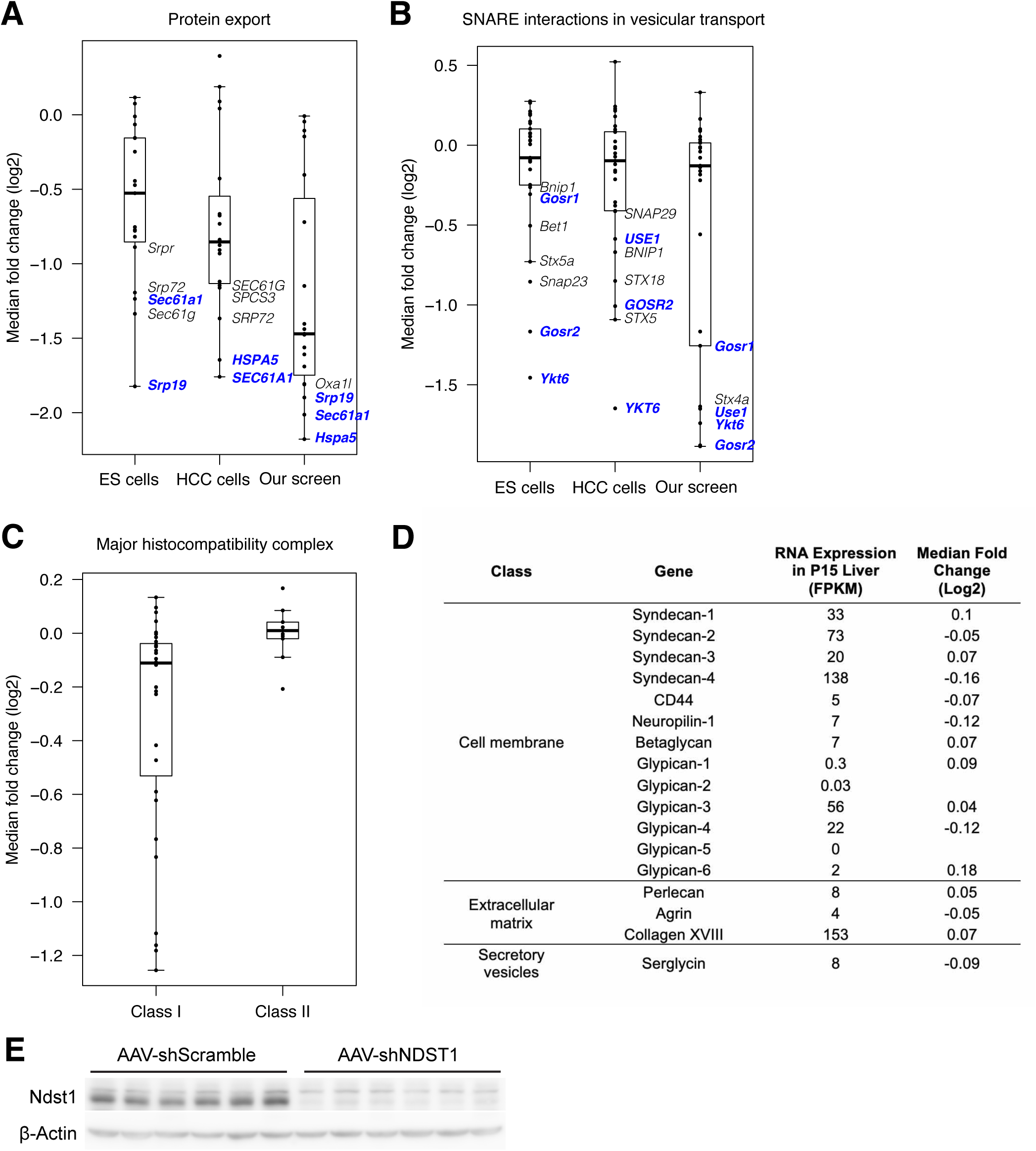
Class I MHC and heparan sulfate biosynthesis are uniquely required for hepatocyte fitness in the organism, Related to Figure 5. (A) Quantile-normalized median fold change (log2) of genes in the KEGG gene set for protein export in ES cell screens, HCC cell line screens, and our screen. Genes depleted in more than one screen are highlighted in blue. The bounds of the box indicate the first and third quartiles, and the whiskers extend to the furthest data point that is within 1.5 times the interquartile range. (B) Quantile-normalized median fold change (log2) for genes in the KEGG gene set for SNARE interactions in vesicular transport in ES cell screens, HCC cell line screens, and our screen. Genes depleted in more than one screen are highlighted in blue. The bounds of the box indicate the first and third quartiles, and the whiskers extend to the furthest data point that is within 1.5 times the interquartile range. (C) Median fold change (log2) across mice for genes associated with class I MHC or class II MHC within the KEGG gene set for antigen processing and presentation in our screen. The bounds of the box indicate the first and third quartiles, and the whiskers extend to the furthest data point that is within 1.5 times the interquartile range. (D) The RNA expression (FPKM) in postnatal day 15 liver and median fold change (log2) across mice in our screen for each gene in the three classes of heparan sulfate proteoglycans. (E) Western blot showing NDST1 protein levels in postnatal day 15 livers injected with 4 x 10^11^ GC of either AAV-shScramble or AAV-shNDST1 on postnatal day 5. Beta-actin is shown as a loading control. The first three lanes for each condition represent livers from male mice while the last three lanes for each condition represent livers from female mice.

**Supplemental Table 1. RNA sequencing data, Related to Figure 3.** Average RNA expression (FPKM) for genes expressed in liver during development, adulthood, and regeneration. n = 3 male mice per time point.

**Supplemental Table 2. sgRNA library, Related to Figure 3.** Sequences for all sgRNAs used in our library and list of control sgRNAs used in our library, Shohat et al. 2019, and Tzelepis et al. 2016.

**Supplemental Table 3. Screen results, Related to Figures 3-5.** Count data for all sgRNAs in the plasmid library, induction library, and individual endpoint libraries, gene scores across mice with test for depletion, gene scores across mice with test for enrichment, list of control genes used to validate screen.

**Supplementary Table 4. Gene set enrichment analysis, Related to** **Figure 5**. Depleted gene sets in our screen, depleted gene sets in our screen versus mouse embryonic stem (ES) cell screens, depleted gene sets in our screen versus human hepatocellular carcinoma (HCC) cell line screens, and gene scores for genes in indicated gene sets in our screen, mouse ES cell screens, and human HCC cell line screens

